# The dynamical Matryoshka model: 1. Incoherent neutron scattering functions for lipid dynamics in bilayers

**DOI:** 10.1101/2021.09.21.461198

**Authors:** Dominique J. Bicout, Aline Cisse, Tatsuhito Matsuo, Judith Peters

**Affiliations:** Univ. Grenoble Alpes, CNRS, Grenoble INP, VetAgro Sup, TIMC, 38000 Grenoble, France; Univ. Grenoble Alpes, CNRS, LiPhy, Grenoble, France; Institute for Quantum Life Science, NIQRST, Tokai, Ibaraki, 319-1106, Japan; Institut Laue-Langevin, 71 Avenue des Martyrs, 38042 Grenoble, France; Institut Universitaire de France, France

**Keywords:** lipids, bilayers, local dynamics, quasi-elastic neutron scattering, modeling

## Abstract

Fluid lipid bilayers are the building blocks of biological membranes. Although there is a large amount of experimental data using inconsistent quasi-elastic neutron scattering (QENS) techniques to study membranes, very little theoretical works have been developed to study the local dynamics of membranes. The main objective of this work is to build a theoretical framework to study and describe the local dynamics of lipids and derive analytical expressions of inconsistent diffusion functions (ISF) for QENS. As results, we developed the dynamical Matryoshka model which describes the local dynamics of lipid molecules in membrane layers as a nested hierarchical convolution of three motional processes: (i) individual motions described by the vibrational motions of H-atoms; (ii) internal motions including movements of the lipid backbone, head groups and tails, and (iii) molecule movements of the lipid molecule as a whole. The analytical expressions of the ISF associated with these movements are all derived. For use in analyzing the QENS experimental data, we also derived an analytical expression for the aggregate ISF of the Matryoshka model which involves an elastic term plus three inelastic terms of well-separated time scales and whose amplitudes and rates are functions of the lipid motions. And as an illustrative application, we used the aggregated ISF to analyze the experimental QENS data on a lipid sample of multilamellar bilayers of DMPC (1,2-dimyristoyl-*sn*-glycero-3-phosphocholine). It is clear from this analysis that the dynamical Matryoshka model describes very well the experimental data and allow extracting the dynamical parameters of the studied system.

## 1. Introduction

Biological membranes are complex lipid-rich systems that constitute fundamental interfaces and selectively permeable barriers for the compartmentalization that defines cells and organisms. Biological membranes are composed of lipids, which self-assemble into bilayers, proteins and carbohydrates [1]. In addition to separating the interior from the exterior of cells, for example, membranes are very dynamic systems that host and ensure many essential processes vital for cellular functions, such as the transport of proteins or ions [1, 2]. This dynamics is made up of both the activity at the surface and the movements of the bilayers which give the membrane an important fluid character to ensure its functions. Since lipids are the most abundant constituents of membranes, studies on the dynamics of lipids in membranes are and remain very crucial. Due to their complex structure and dynamics, lipid bilayers are characterized by a hierarchy and heterogeneity of motions over a wide range of time and space. These dynamics include, for example, local movements like lipid rotational, in-and-out of the plane diffusion at very short spatial scales and time scales of pico to nanoseconds, but also collective lipid movements like density fluctuations of short wavelength in pico to nanosecond range and long-wavelength flip-flop or undulation and bending modes of the bilayer in the nano to microsecond range [2, 3, 4]. In this work, we will only deal with short-range movements carried over short time scales (pico to nanoseconds).

Over the years, incoherent quasi-elastic neutron scattering (QENS) has proven to be a key technique for investigations of lipid motions at the pico - to nanosecond time scale and there is a great amount of experimental data that have been accumulated using QENS techniques [5, 6, 7, 8, 9, 10]. However, since the seminal work by Pfeiffer et al. [5], there are very few theoretical works that have been developed to analyze and describe local dynamics of membranes [5, 11]. Along these lines, a study on phospholipid membranes has been proposed to separate the motions over three distinct time scales [12]. Using such models with QENS data allow to retrieve parameters like mean size of the solvent lipid cages, diffusion coefficients lipid rotations or in-plane Brownian motions. Motivated by these experimental investigations and findings, our main objective in this paper is to provide a theoretical framework of a model of membrane layers and to derive analytical expressions of incoherent structure functions (ISF) describing the local dynamics. To this end, we have developed a model that describes a membrane layer as a system of dynamically equivalent lipid molecules, each of which consisting of two connected (via a backbone) bodies (head and tail) undergoing kinds of internal and body motions.

The remaining of the paper is organized as follows. Section 2 consists of three parts: (i) formulation of the dynamical Matryoshka model which describes the hierarchical convolution of movements in the dynamics of a lipid molecule, (ii) detailed description of motions that are included in the model and derivations of analytical expressions of the associated ISFs, and (iii) derivation of an aggregate expression of the global ISF to be used to analyze and fit the experimental data. Technical details of all derivations are described in the Appendices A - C. Section 3 illustrates how the developed theory can be used to analyze the experimental QENS data of a bilayer membrane and extract the parameters of interest. Finally, the main results of the paper are summarized in the Section 4.

## 2. Model formulation and analytical expressions

The membrane layer is considered as a system made of structurally and dynamically equivalent lipid molecules interacting with each other. Among the whole hierarchy of motions characterizing the membrane dynamics, we are interested in the local motions of lipid molecules as can be studied using techniques like incoherent quasi-elastic neutron scattering, inelastic x-ray scattering, to cite a few [5, 6, 7, 8, 9, 10]. Figure 1A provides an illustration of the types of phospholipid molecules we will be dealing with. To describe the motional processes of an individual lipid molecule, occurring in the potential of mean force generated by the sea of lipid molecules in the membrane layer, we develop the Matryoshka model described below (see Fig. 1B).

**Figure 1:**
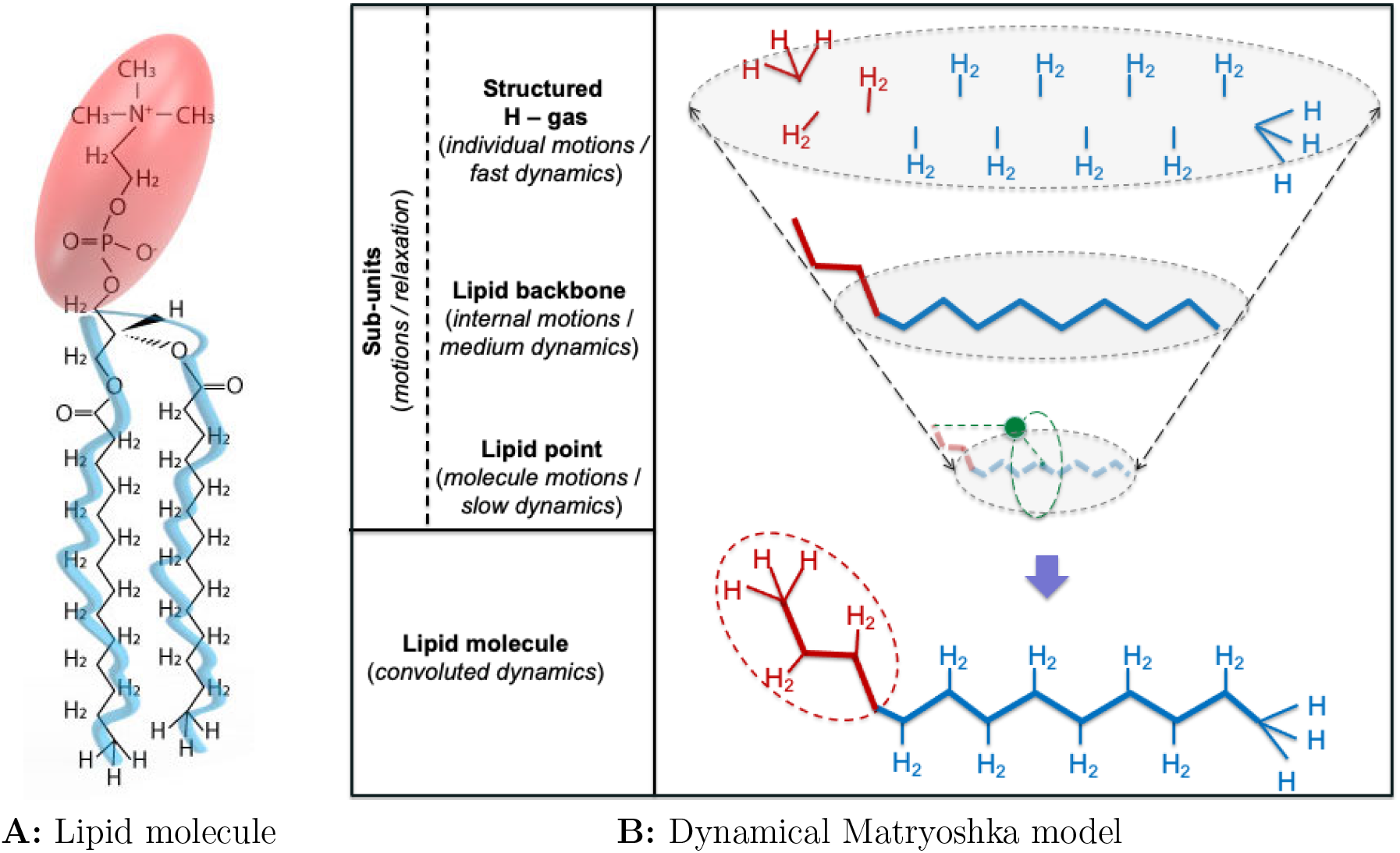
**A:** Illustration of a phospholipid molecule 1,2-dimyristoyl-*sn*-glycero-3-phosphocholine (DMPC) consisting of a hydrophilic head (in red) and hydrophobic fatty acid tails (in blue) with the distribution of H-atoms. Courtesy from Steph Monfront (ILL). **B: Dynamical Matryoshka model** for the dynamics of the lipid molecule represented as a funnel of three - level convoluted dynamic processes. Lipid tails are represented by an effective single tail for the dynamics. Red and blue colored elements relate to head and tail groups, respectively. H-atoms with C-H bounds (top) are structured along the lipid backbone (middle), with head and tail subunits, mirroring the lipid molecule representation (bottom of the figure). At the bottom of the funnel, the lipid molecule as a whole (rigid body) comes down to a point particle (green sphere) representing the center of inertia (barycenter) of all dynamic H-atoms with respect to the lipid main axis (dashed lipid backbone).

### 2.1. Dynamical Matryoshka model

As illustrated in Fig. 1B, the dynamical Matryoshka model describes the dynamics of a lipid molecule as resulting from the hierarchical convolution of three motional processes (from fastest to slowest motions): (i) *individual motions* of H-atoms forming the cloud of structured H-atoms bound to backbone atoms, (ii) *internal motions* of the lipid backbone or skeleton made up of non-H atoms and H-atoms bound to them, the head and tail subunits, and (iii) *molecule motions* or rigid body motions of the lipid molecule as a whole represented by the movements of the center of inertia all dynamic H-atoms with respect to the lipid main axis. The metaphor of dynamical Matryoshka (nesting dolls) originates from what these motional processes occur in a superimposed and nested way over various (and overlapping) timescales and, therefore, are resolved by zooming in or out over the associated characteristic timescales.

### 2.2. Dynamical Matryoshka model: Incoherent Structure Function (ISF)

Within the framework of this model and the hypothesis of dynamic independence for different motions, the ISF for the local motions of a lipid molecule can be written as,

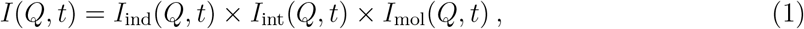

where *I*_ind_(*Q, t*), *I*_int_(*Q, t*) and *I*_mol_(*Q, t*) are the ISFs associated to individual, internal and molecular motions, respectively. In the following, we detail the motional processes of lipids included in the dynamical Matryoshka model with, whenever possible, the associated potentials of mean force, *V* (**r**). The elastic incoherent structure factor (EISF) for each motion can be calculated as (see, Table 4),

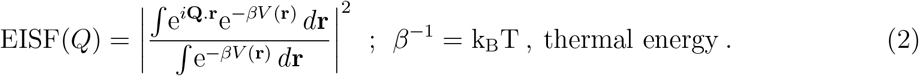

The relationships between all the dynamical processes are summarized in Table 1 with the details of derivations of IFSs provided in Appendix A and the analytical expressions of the ISFs reported in Table A.7.

**Table 1:** Dynamical processes.

| Dynamics | Subunit | Motions | Process |
| --- | --- | --- | --- |
| Individual | H-atoms | *vibrational Debye-Waller* | Very Fast |
| Internal | C-H groups (backbone) $\otimes$ Tail | *jump dynamics* $\otimes$ *diffusion in circles* | Fast |
| | Head | *rotational diffusion* $\otimes$ *flip-flop* | Intermediate |
| Molecule | Lipid Molecule | $\otimes$<br>*rotational diffusion* | |
|  |  | *in-out of the plane diffusion* | Slow |
| | | $\otimes$<br>*2d-diffusion in a cage* | |

#### 2.2.1. Individual motions

Individual motions relate to the vibrational motions of the cloud of H-atoms bound to the lipid backbone atoms. Such motions are described by a harmonic potential of mean force, *V* (*r*) = *r*^2^*/*2 ⟨*u*^2^⟩, where ⟨*u*^2^⟩ is the ensemble mean-square displacements of H-atoms about their equilibrium positions. These individual movements being relatively fast (∼ 100 meV, [13]) for the timescales that concern us, the associated ISF is reduced to the EISF (i.e., time independent) and, in fact, is factored out in Eq.(1). The ISF for individual motions is therefore given by the Debye-Waller factor as, *I*_ind_(*Q, t*) = *I*_DW_(*Q*) = *A*_DW_(*Q*) (see Table A.7).

#### 2.2.2. Internal motions

Internal motions result from the combination of three motional processes from the lipid backbone, head and tail subunits. The headgroup and tails are considered as independent subunits each with *z* and 1 − *z* fraction of H-atoms, respectively, and the lipid backbone common to both subunits includes all H-atoms (see Fig. 2). The ISF for internal motions writes as,

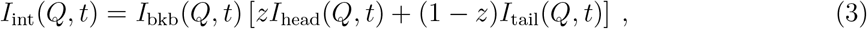

where *I*_bkb_(*Q, t*), *I*_head_(*Q, t*) and *I*_tail_(*Q, t*) are the ISFs for motions in the lipid backbone, head and tail.

**Figure 2:**
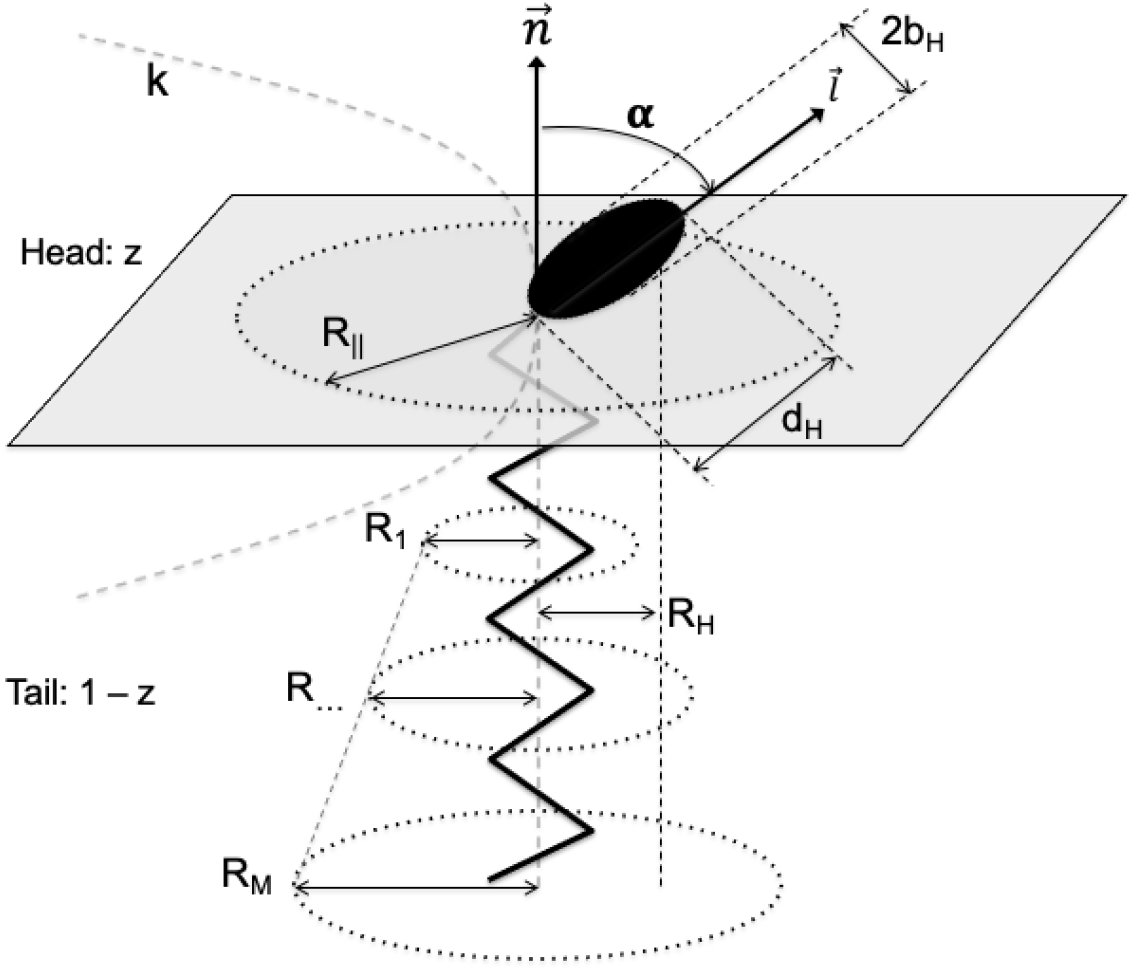
Scheme of a lipid molecule showing the characteristic parameters of motions. The polar head group is represented by the black ellipse (tilted by an angle *α*) of short and long diameters 2*b*_H_ and *d*_H_, respectively, and the hydrophobic tails are represented by a dynamically effective tail (vertical zigzag black line) of apparent length *M*. Grey plane separates the head group (with a fraction *z* of H-atoms) from the tail (with a fraction 1 − *z* of H-atoms). The membrane normal (lipid axis) and head axis (tilted by an angle *α*) are indicated by 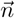 and 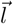, respectively. Dashed circle of radius *R*_||_ in the grey plane represents the effective cage for the in-plane diffusion and the dashed parabola of stiffness *k*, perpendicular to the membrane plane, indicates in-out of the plane motions of the lipid molecule. Dashed circles of increasing radii, *R*_1_, …, *R*_*M*_, around the effective tail represent areas for the diffusion motions of Hs along the tails and *R*_H_ is the distance between the lipid axis 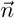 and the center of inertia of all H-atoms with respect to 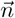.

- *ISF for the Backbone motions, I*_bkb_(*Q, t*): Backbone motions relate to the motions of C-H groups in the lipid molecule. Several models can be used to describe the heterogeneous dynamics of C-H groups depending on whether they are located in the head group or in the tails. For simplicity, we considered that the movements of all the C-H groups could be described using a jump dynamics between two non-equivalent sites distant from *d* and characteristic relaxation rate Γ_jump_. Because of heterogeneity, *d* and Γ_jump_ are ensemble averaged quantities over distributions of site distances and relaxation times, respectively. The potential of mean force for these motions is defined as [14],

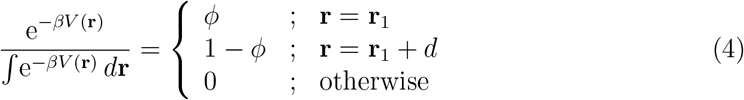

where **r**_1_ is the position of the low energy site and *ϕ* is the probability of occupying **r**_1_ (see Table A.7) and the ISF, *I*_bkb_(*Q, t*) = *I*_jump_(*Q, t*), is given in Table A.7.
- *ISF for Tail motions, I*_tail_(*Q, t*): Tail motions refer to the movements of the H-atoms in all the tails. In Fig. 2, the tails, which interact with each other, are represented by a dynamically effective tail which indicates the axis of the tails and especially the surfaces within which the movements of the H-atoms take place. The movements of H-atoms in the tails are modeled by classical two-dimensional diffusions, all of an effective diffusion constant *D*_tail_ within circles of radii *R*_*m*_ in the plane parallel to the membrane at *m*, where *m* designates the position of the carbon atoms *C* − *C* along the lipid tail from the start to the tail end (with, 1 ≤ *m* ≤ *M*) and *M* is the apparent length of the lipid tail. As shown in Fig. 2, as a result of interactions between tails, *R*_*m*_ increases with *m* as the explorable space increases with *m*. Unlike the linear increase of Carpentier et al. [15], we use instead, 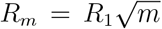, to somehow account for a random walk-like (along the tail axis) positions of carbon atoms to which hydrogens are bound. The potential of mean force for these motions at each *m* is,

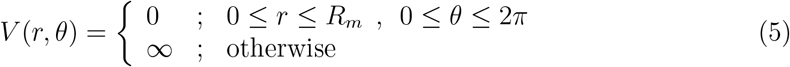

and the ISF *I*_tail_(*Q, t*) is given in Table A.7.
- *ISF for Head motions, I*_head_(*Q, t*): Headgroup motions, involved in the roughness of the membrane surface, include three independent motions (see Fig. 2): the uniaxial rotational diffusion about the head axis 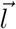, the flip-flop or jump dynamics of 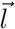 axis between angles −*α* and *α* about the membrane normal 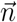 axis and the uniaxial rotational diffusion of the 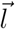 axis about the membrane normal 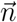 axis. However, for our purpose, by symmetry the rotational diffusion about the 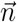 axis averages out such that only the rotation about 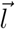 and the flip-flop about 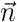 are retained. The headgroup size *b*_H_ (see Fig. 2) represents the distance between the head axis and the center of inertia of all H-atoms of the head group with respect to the head axis 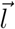. Derivation of the ISF *I*_head_(*Q, t*) is detailed in Appendix A.

#### 2.2.3. ISF for molecule motions: I_mol_(Q, t)

For molecule or rigid body movements, all dynamic H-atoms together perform the same processes regardless of their location in the lipid molecule. Formally, the dynamics of the lipid molecule as a whole can therefore be described by that of the center of inertia of all dynamic H-atoms with respect to the lipid main axis (Fig. 1B). Three types of independent movements are considered: rotational motions around the axis 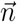 of the lipid molecule, in-out of the membrane plane movements (parallel to 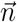) and in-plane lateral diffusion of the lipid molecule. The ISF for molecular motions writes as,

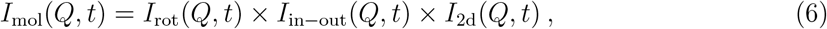

where *I*_rot_(*Q, t*), *I*_in*−*out_(*Q, t*) and *I*_2d_(*Q, t*) are the ISFs for rotational, in-out of the plane and in-plane diffusion motions.

- *ISF for rotational motions, I*_rot_(*Q, t*): For the rotational motions of the lipid molecule about its axis 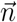, all H-atoms perform exactly the same rotational movement about the lipid axis 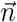. However, as not all of the H-atoms are located equidistant from the lipid axis, the resulting ISF should be the sum of the individual ISFs. As a first approximation, we can reduce the rotational motions of all H-atoms to that of their center of inertia and describe the rotational motions of the lipid molecule about its axis 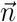 by a rotational diffusion of diffusion coefficient *D*_rot_ on a circle of radius *R*_H_, where *R*_H_ is the distance between the lipid axis and the center of inertia of all dynamic H-atoms with respect to the lipid axis 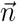. The ISF *I*_rot_(*Q, t*) is given in Table A.7.
- *ISF for in-out of the plane motions, I*_in*−*out_(*Q, t*): The in-out-of-plane movements, involved in the roughness of the membrane surface, relate on the up and down motions (normal to the membrane plane) around the equilibrium position of the lipid molecule within the membrane layer. Such motions can be described by the one-dimensional (parallel to 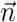) diffusion within a harmonic potential of force constant *k* and relaxation time *τ*. The harmonic potential of mean force is given by, *V* (*z*) = *kz*^2^*/*2, where *z* is the lipid molecule position around its equilibrium position. The ISF *I*_in*−*out_(*Q, t*) is given in Table A.7.
- *ISF for in-plane 2d diffusion motions, I*_2d_(*Q, t*): In general, in-plane lateral diffusion of lipid molecules is composed of short range local diffusion in a solvent cage and long range jumps between different sites allowing molecules to travel within the membrane layer. As this work is dealing with spatially short range dynamics, we will only consider short range diffusion in what follows. Thus, in-plane diffusion of lipid molecules describes when molecules exchange places via Brownian motion within a cage. Such motions are described by a two-dimensional diffusion of diffusion constant *D*_||_ within a circle of radius *R*_||_, where *R*_||_ is the effective size of the cage formed by the neighboring lipid molecules. The potential of mean force for such a confined isotropic two-dimensional diffusion is given by,

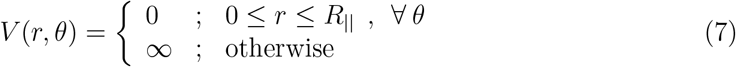

where *r* is the position of lipid in the membrane plane. The ISF *I*_2d_(*Q, t*) is given in Table A.7.

All the 7 movements included in the dynamical Matryoshka model and associated 18 dynamical parameters are summarized in the Table 2.

**Table 2:**
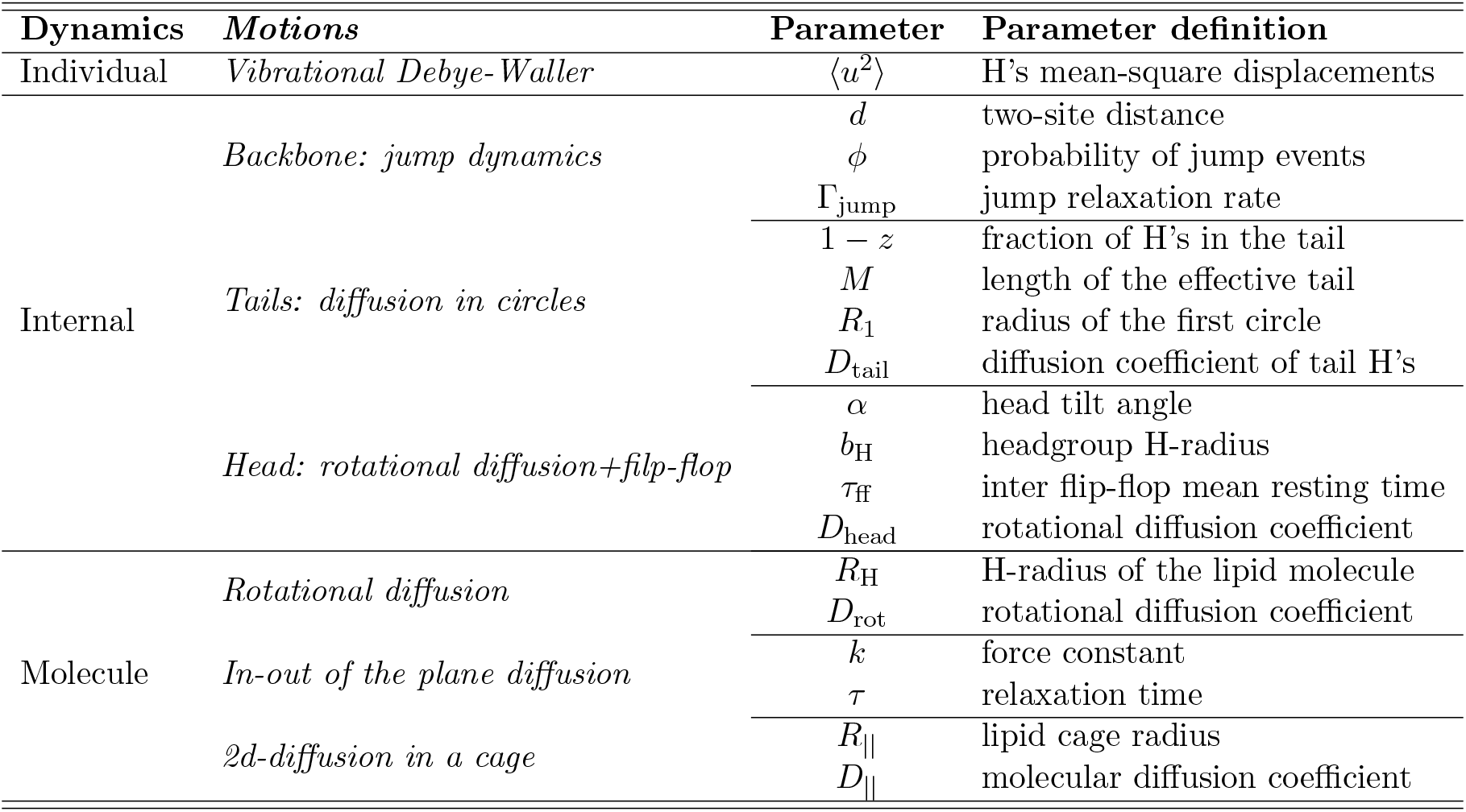
Local motions and associated parameters of the Matryoshka model.

### 2.3. Aggregated processes: 3-dynamical process approximation

As outlined above, the ISF of local movements of lipid molecules results from the convolution of several individual ISFs which, in general, are multi-exponential functions of time. Even when each individual ISF was approximated by a single exponential relaxation, the resulting ISF would still be multi-exponential. In practice, it would be challenging to use such a multi-exponential function in the analyzes of experimental data to extract physical parameters of interest listed in Table 2. The idea is therefore to develop an approximation of the ISF allowing to reduce timescales by coupling them together while keeping the original information (as summarized in Table 2). For that purpose, we follow Wanderlingh et al. [12] and consider how the dynamics of the lipid molecules as described above can be approximatively aggregated into 3-dynamical processes. Under the context of dynamical processes taking place on widely separated time scales (one or few orders of magnitude) as outlined above, the main value of the 3-dynamical process model is to allow fitting the experimental data and, therefore, providing an opportunity for direct connection between analytical models of lipid motions and experimental observations.

The 3-dynamical process model is a phenomenological model in which the position **r**(*t*) at time *t* of the H-atom in lipid molecules can be split into three independent components, **r**(*t*) = **r**_1_(*t*) + **r**_2_(*t*) + **r**_3_(*t*), where **r**_1_(*t*), **r**_2_(*t*) and **r**_2_(*t*) stand for the slow, intermediate and fast motions, respectively, describing the overall motions of hydrogen atoms in the equilibrium lipid molecule at very different time scales. Each of these motions accounts for a confined dynamics with a quasi-elastic term described by a single exponential decay term. Under the approximation of separate time scales, the aggregated ISF can be written as,

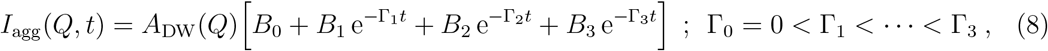

where *A*_DW_(*Q*) stands for the Debye-Waller factor, *B*_0_ is the overall EISF, *B*_*i*_ and Γ_*i*_ for *i* = 1, 2, 3 are the amplitudes and relaxation rates of the slow, intermediate and fast motions, respectively, such that, 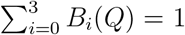 for all *Q*. Accordingly, the measurable function in incoherent neutron scattering experiments is,

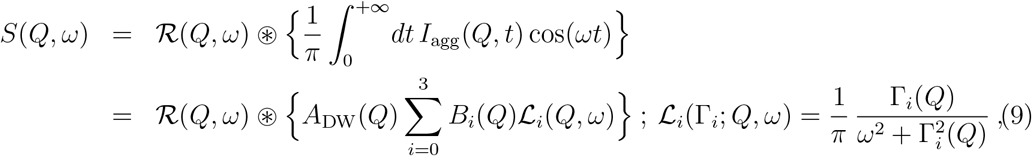

where ℛ(*Q, ω*) refers to the experimental resolution function, *B*_*i*_(*Q*) and Γ_*i*_(*Q*) are the respective areas and half-widths at half-maximum (HWHM) of the Lorentzian functions ℒ_*n*_(*Q, ω*). It should be stressed here that:

i. the three time scales slow, intermediate and fast do not necessarily correspond or coincide to and, therefore, are not to be confused with those of individual, internal and collective motions (see Table 1);
ii. the phenomenological model does not provide which movements are resolved at each timescale.

The purpose of aggregating the *I*(*Q, t*), derived above and given in Eq.(1), into 3-dynamical processes is to associate each term in Eq.(8) with a physical meaning and an analytical expression. To this end, we use the mapping, *I*(*Q, t*) ≈ *I*_agg_(*Q, t*), i.e.,

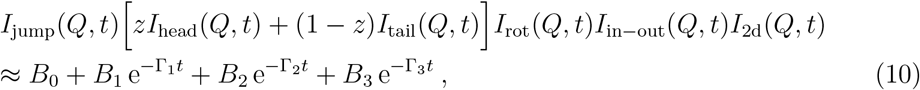

to derive the expressions of *B*’s and Γ’s as functions of physical quantities in *I*(*Q, t*). Note that in Eq.(10), *B*_0_ is already known and is equal to *B*_0_ = *I*(*Q, t* → ∞) = *I*_agg_(*Q, t* → ∞) (i.e., the product of EISF of all local motions), whereas the *B*_*i>*0_ and Γ_*i>*0_ remain to be determined. In addition, the number of terms after the approximate sign “≈” in Eq.(10) can be reduced when some of the amplitudes cancel out because the contributing motions turn out to be not observable. The aggregation procedure, based on the hierarchy of relaxation time scales of motional processes in the Matryoshka model, is detailed in Appendix B and the results are summarized in Table 3 in which the expressions of original EISFs and relaxation rates are given in Tables 4 and 5, respectively. Table 3 provides which local motions contribute at each timescale of the 3-dynamical process and how they are combined in the expressions of amplitudes and relaxation rates. Each relaxation rate is not exclusive of a single local motion and the same local motion can therefore contribute to different timescales. The amplitudes result from the combination of several local motions whose relaxation times can be relatively different. From the amplitudes (with the Debye-Waller factoring all), we have the following contributions:

**Table 3:** Amplitudes (areas) and relaxation rates of the 3-dynamical process approximation. 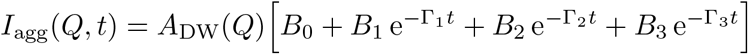.

| Motions | Amplitudes (areas) |  |
| --- | --- | --- |
|  | theoretical<sup>(a)</sup> | experimental<sup>(b)</sup> |
| EISF | $B_0 = [zA_{\text{head}} + (1 - z)A_{\text{tail}}] A_{\text{jump}}A_{\text{rot}}A_{\text{in-out}}A_{2\text{d}}$ | $A_0 = mB_0 + \varepsilon_0$ |
| Slow | $B_1 = [zA_{\text{head}} + (1 - z)A_{\text{tail}}] (1 - A_{\text{in-out}}A_{2\text{d}})A_{\text{jump}}A_{\text{rot}}$ | $\left. \vphantom{\begin{matrix} B_0 \\ B_1 \\ B_2 \end{matrix}} \right\} A_i = mB_i + \frac{[1 - m - \varepsilon_0]}{3}$ |
| Intermediate | $B_2 = A_{\text{jump}} [z(1 - A_{\text{head}}A_{\text{rot}}) + (1 - z)A_{\text{tail}}(1 - A_{\text{rot}})]$ | |
| Fast | $B_3 = z(1 - A_{\text{jump}}) + (1 - z)(1 - A_{\text{jump}}A_{\text{tail}})$ | |
|  | Relaxation rates |  |
| Slow | $\Gamma_1 = \frac{(1 - A_{\text{in-out}})\Gamma_{\text{in-out}} + (1 - A_{2\text{d}})\Gamma_{2\text{d}}}{1 - A_{\text{in-out}}A_{2\text{d}}}$ | |
| Intermediate | $\Gamma_2 = \frac{z(1 - A_{\text{head}})\Gamma_{\text{head}} + [z + (1 - z)A_{\text{tail}}] (1 - A_{\text{rot}})\Gamma_{\text{rot}}}{z(1 - A_{\text{head}}A_{\text{rot}}) + (1 - z)A_{\text{tail}}(1 - A_{\text{rot}})}$ | |
| Fast | $\Gamma_3 = \frac{(1 - A_{\text{jump}})\Gamma_{\text{jump}} + (1 - z)(1 - A_{\text{tail}})\Gamma_{\text{tail}}}{z(1 - A_{\text{jump}}) + (1 - z)(1 - A_{\text{jump}}A_{\text{tail}})}$ | |

<sup>(a)</sup> Limiting properties for $B_i$ : $B_0(Q \rightarrow 0) = 1$ and $B_{i>0}(Q \rightarrow 0) = 0$ ; $B_i(\text{high } Q) \rightarrow 0$ for both $i = 0$ and $i = 1$ while $B_2(\text{large } Q) = 1 - B_3(\text{large } Q) = zA_{\text{jump}}(\text{large } Q)$ where $A_{\text{jump}}(\text{large } Q) = A_{\text{bkb}}(\text{large } Q) \approx 1 - 2\phi(1 - \phi)$ .
<sup>(b)</sup> See Appendix C for derivations. $m$ is the fraction of dynamical or mobile H-atoms and the error function $\varepsilon_0(Q)$ accounts for immobile H-atoms, multiple scattering effects observable at low- $Q$ , and others. For simplicity, we will assume that the errors are homogeneously distributed over length scales and, therefore, $\varepsilon_0(Q) = \varepsilon_0$ .

**Table 4:**
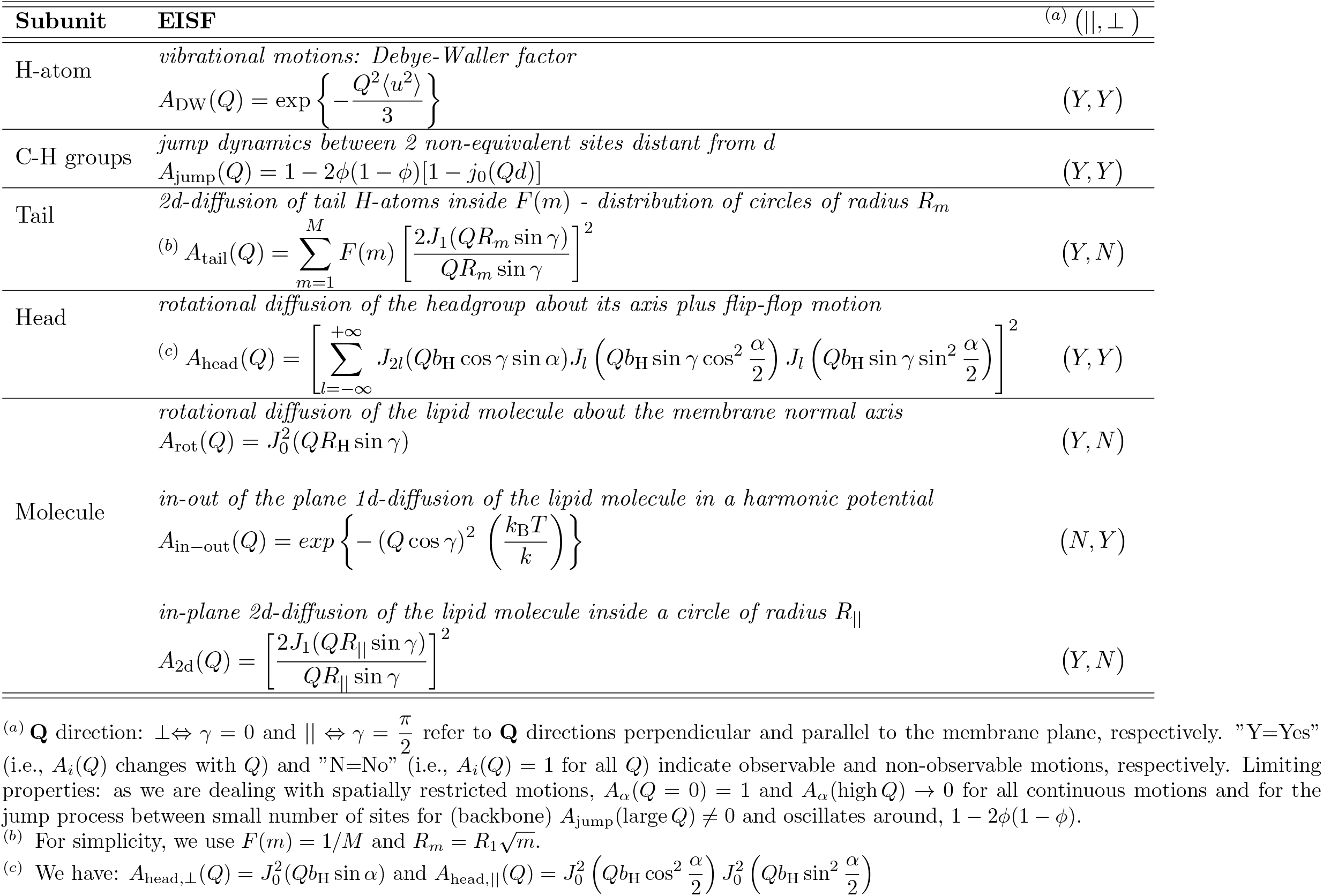
Elastic Incoherent Structure Factor (EISF) of dynamical processes.

**Table 5:**
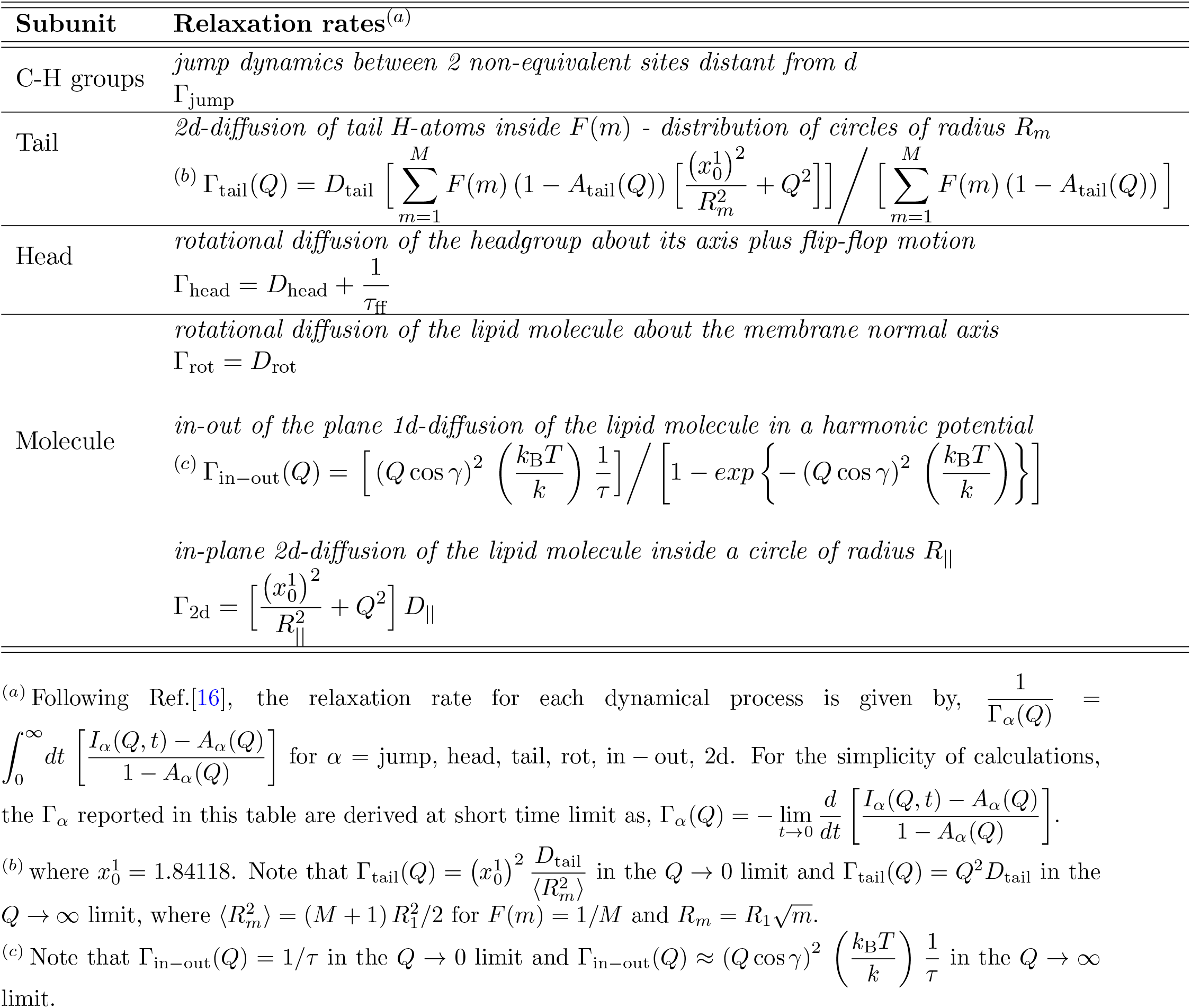
Relaxation rates of dynamical processes.

- *Slow motions:* backbone, head, tails, rotational, in-out of the plane and in-plane diffusion;
- *Intermediate motions:* backbone, head, tails and rotational;
- *Fast motions:* backbone and tails.

And, as a function of *Q*, we have:

- *Q* → 0 limit: as expected, the scattering function essentially consists of the elastic peak (EISF) with all motions contributing and no Lorentzian functions as, *B*_0_(*Q* → 0) → 1 and *B*_*i>*0_(*Q* → 0) → 0;
- Moderate *Q*: as all *B*_*i*_ ≠ 0, the scattering function consists of the elastic peak plus the three Lorentzian functions with contributions of all motions. Note that slow motions are only observable for these *Q*-ranges as *B*_1_(*Q*) → 0 in both *Q* → 0 and large *Q* limits;
- Large *Q* limit: as *B*_*i≤*1_(large *Q*) → 0 and *B*_2_(large *Q*) = 1 − *B*_3_(large *Q*) ≈ *zA*_bkb_(large *Q*) ≠ 0, the scattering function reduces to only two Lorentzian functions involving intermediate and fast motions and concerning motions of C-H groups of the backbone.

## 3. Illustrative Example

To compare and test how the theoretical developments outlined above would work when analyzing experimental data, we considered the following neutron scattering experiments. The whole task consists in determining 18 parameters describing the 7 motions of H-atoms in the Matryoshka model (see, Table 2) plus 2 experimental parameters (*m* and *ε*_0_ in Table 3).

### 3.1. Quasi-Elastic Neutron Scattering (QENS) experiments and analyses

We used lipid samples of multilamellar bilayers (MLBs) of DMPC (1,2-dimyristoyl-*sn*-glycero-3-phosphocholine) represented in Fig. 1A. The DMPC was purchased from Lipoid (Ludwigshafen Germany) or from Avanti Polar Lipids (Alabaster, USA) and used without further purification. The lipid samples were prepared on 6 Si wafers, and hydrated in D_2_O atmosphere within a desiccator at full hydration. They were mounted on flat Aluminum sample holders, gold-coated to avoid sample contamination. The sealing of the cells was done by using Indium wire, and they were weighed before and after the experiment to check for any sample loss.

DMPC samples were scanned on the IN6 time-of-flight spectrometer from ILL (Grenoble, France), with a wavelength of 5.1 Å, corresponding to an energy resolution of about 70 µeV [17]. At this resolution, motions up to around 10 ps are accessible and the attainable *Q*-range is of, 0.37 ≤ *Q* ≤ 2.02 *Å*^−1^. The sample holder was oriented at 135° from the beam to access in-plane motions [8]. QENS scans were performed at three different temperatures, 280 K, 311 K and 340 K, to probe the dynamics below and above the main phase transition of lipid bilayers; DMPC is known to undergo consecutive phase transitions from the gel phase to the ripple phase around 287 K and to the fluid phase around 297 K (corresponding to the physiological state in cells) [18].

Empty cell with and without wafers, as well as Vanadium, were measured for correction and normalization purposes. Raw data were first corrected by the empty cell + 6 wafers contribution, using the Large Array Manipulation Program (LAMP) [19]. The resulting S(*Q, ω*) spectra were subsequently analyzed in the range of -10 meV ≤ ΔE ≤ 2 meV using IGOR Pro software (WaveMetrics, Lake Oswego, OR, USA). Following the QENS analysis in [12], the model used to fit the spectra is similar to that in Eq.(9) as,

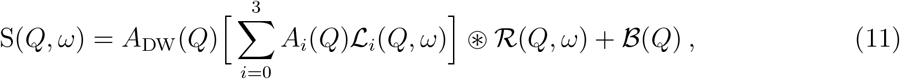

where *A*_0_(*Q*) is the experimental elastic incoherent structure factor (EISF) and the *A*_*i*_(*Q*) for *i >* 0 are the experimental areas of the Lorentzian functions ℒ_*i*_(*Q, ω*) of half-widths at half-maximum Γ_*i*_(*Q*) as defined in Section 2.3. The experimentally determined *A*_*i*_(*Q*) are related to the areas *B*_*i*_(*Q*) in Eqs.(8) and (9) by expressions derived in Appendix C and given in Table 3. ℛ(*Q, ω*) refers to the resolution function and corresponds to the Vanadium measurements directly included in the analysis and ℬ (*Q*) is a flat background, that can include the instrument contribution, or fast vibrational motions outside the window.

The areas retrieved from the QENS analyses were fitted with the *A*’s in Table 3 using the package *lmfit* from Python [20] with Levenberg-Marquardt and Nelder-Mead algorithms. Fitting procedure relies on the simultaneous fitting of the four areas with corresponding functions given in Tables 3 and 4 through a set of shared parameters. In this way, both statistics are improved with more data points included and constraints on parameters are controlled.

### 3.2. Results

Main results within the framework of this analysis can be summarized as follows:

- Data from QENS experiments are analyzed as illustrated in Fig. 3 where the Eq.(11) is used to fit the data points and extract both the experimental Lorentzian amplitudes (areas) *A*_*i*_(*Q*)’s and HWHM Γ_*i*_(*Q*)’s. Figure 3 shows that the 3-dynamical process model describes very well (with residuals ∼ 0) experimental data into an elastic peak (*δ*(*ω*), a Lorentzian with Γ_0_ = 0, and amplitude *A*_0_(*Q*)) plus three well separated Lorentzians (with, Γ_1_ ∼ 0.1 meV, Γ_2_ ∼ 1 meV and Γ_3_ ∼ 10 meV, for slow, intermediate and fast motions, respectively; there is an order of magnitude between the Γ_*i*_’s). Such an analysis is performed for all values of the pair (*Q, T*) considered in the experiment.

**Figure 3:**
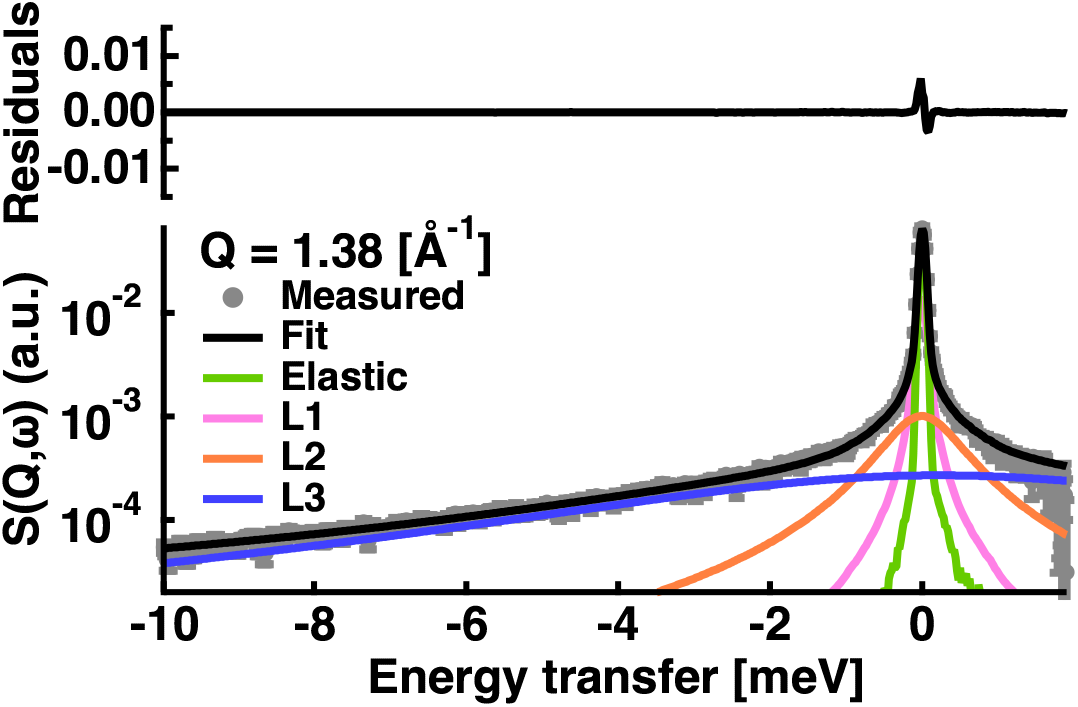
Example of *S*(*Q* = 1.38 *Å*^−1^, *ω*) data fitting for DMPC multilamellar bilayers sample measured on IN6 at 135° geometry (in-plane motions) and T = 280 K. Solid black line through data points (grey circles) represents the best fit to the data using Eq.(11) with the resolution function given by the Vanadium measurements at Q = 1.38 Å^−1^. The fit results from the sum of the elastic peak (green line) and of the three Lorentzian functions for slow (magenta line), intermediate (orange line) and fast (blue line) motions. In the rest of this illustrative example, we will only deal with the *A*_*i*_(*Q*)’s for a single sample in a given geometry whereas the use of this approach on several samples of lipid systems in different geometries with the analysis including *S*(*Q, ω*), *A*_*i*_(*Q*) and Γ_*i*_(*Q*) will be presented and discussed in more details elsewhere [21, 22]. Therefore, the number of parameters to determine is reduced to 11 for the Matryoshka model plus the 2 experimental parameters (*m* and *ε*_0_). All the parameters are listed in the Table 6 where some are fixed (as extracted from the literature) and others are obtained from the best fit of the model to experimental data.

**Table 6:**
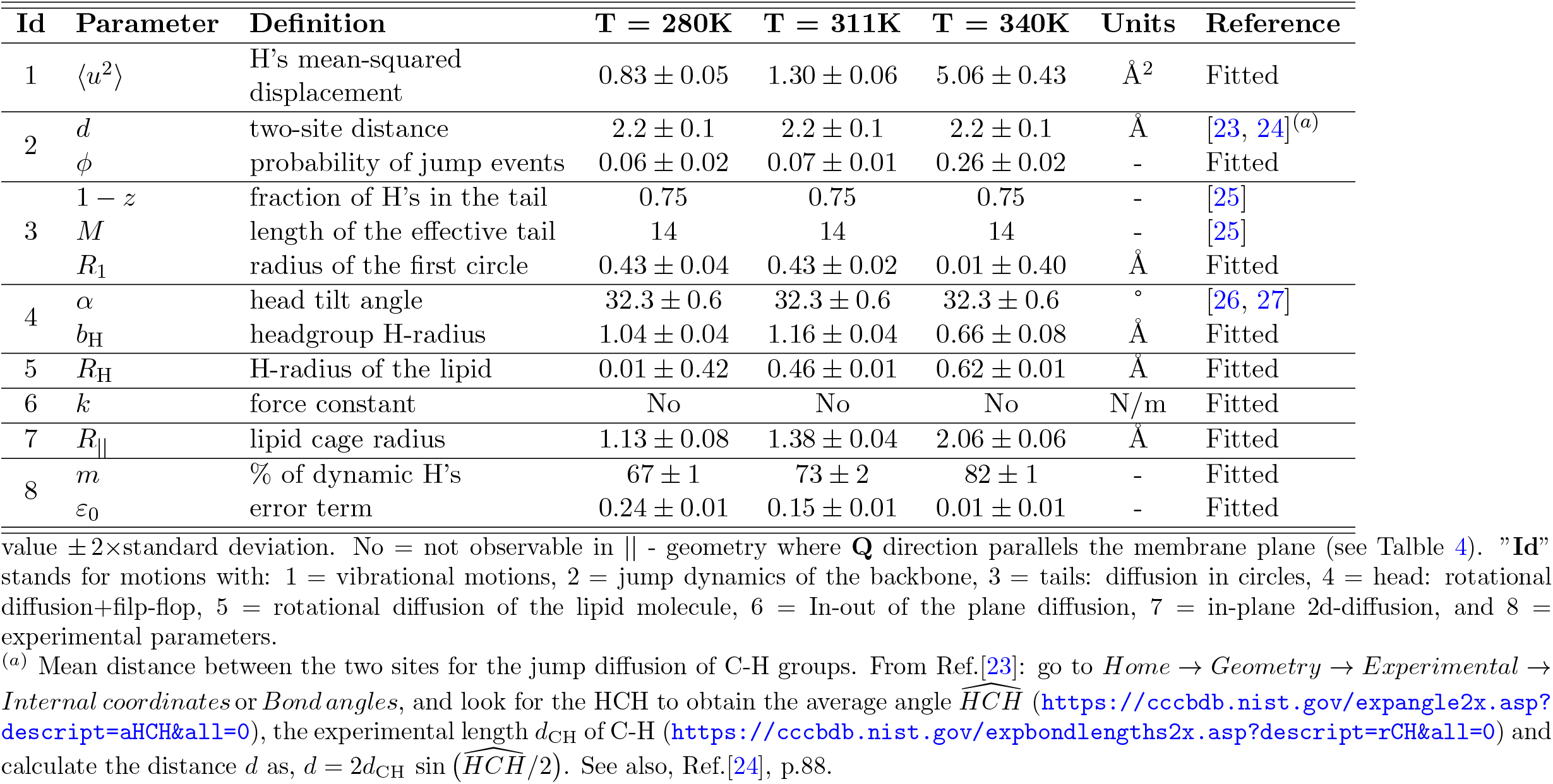
Fixed and Fitted parameters for DMPC MLBs. The sample was measured on IN6 instrument at 135°(in-plane motions or || - geometry).
- Experimental Lorentzian amplitudes (areas) *A*_*i*_(*Q*)’s extracted as described above in Fig. 3 are represented by data points in Fig. 4. We observe that the *A*_*i*_(*Q*) exhibit as a function of *Q* a behavior exactly as predicted by the model (see the end of the Section 2.3): *A*_0_(*Q*) ∼ 1 while *A*_*i>*0_(*Q*) ∼ 0 at low *Q* → 0 as expected for spatially restricted motions, *A*_0_(*Q*) and *A*_1_(*Q*) decreases to very low value (closed to zero) in the large *Q* limit like for continuous motions while *A*_2_(*Q*) and *A*_3_(*Q*) increase with *Q* to a non-zero value reflecting the spatially discrete motions of the backbone.

**Figure 4:**
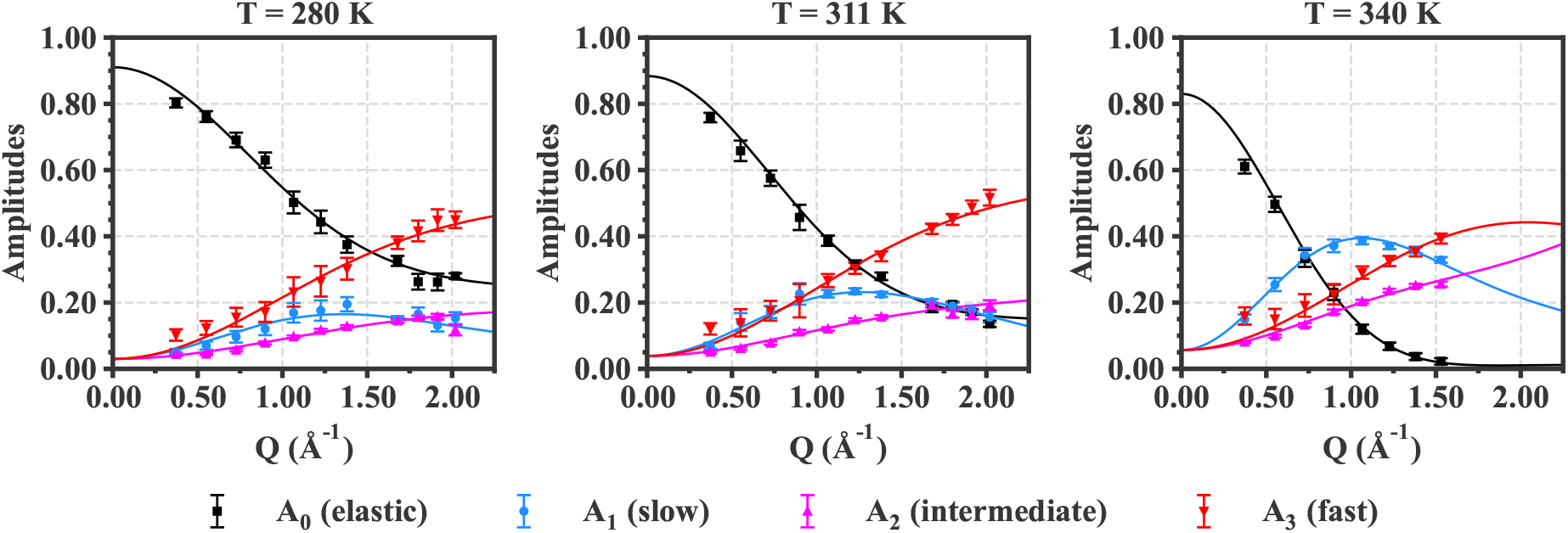
Data fitting for DMPC multilamellar bilayers measured at 135° (in-plane motions). Data points correspond to experimental amplitudes (areas) of the 3-dynamical process as a function of *Q* for different temperatures, obtained from best fits of QENS data with Eq.(11) as illustrated in Figure 3. Solid lines through the data represent best fits to the data using the expressions of amplitudes given in Tables 3 and 4. Parameters extracted from these fits are reported in Table 6. To go further in the analysis of these data, expressions of the amplitudes as a function of Q are needed; this is what we have done in deriving the expressions given in Tables 3 and 4. Lines through the data in Fig. 4 represent best fits (with *χ*^2^ ∼ 1.5) of *A*_*i*_(*Q*)’s using expressions in Tables 3 and 4 with the physical parameters of the local motions thus determined and reported in Table 6. This figure clearly demonstrates the relationships between local motions and the amplitudes of the phenomenological 3-dynamical process model.
- From now on, we will be able to take an interest in local motions. Figure 5 shows the EISF’s or amplitudes (except the Debye-Waller factor) of all local motions contributing to the 3-dynamical process. These EISF’s are generated by using parameters determined from Fig. 4 into expressions in Table 4. Changes with temperature of the variations in amplitudes as a function of *Q* reflect changes of parameters with temperature (see Table 6). Here, we did not nor have developed models to predict how these parameters might change with temperature. We can already notice that in general all parameters exhibit an increase with temperature (see Table 6). Indeed, the lipid system gains thermal energy as temperature increases leading to a subsequent enhancement of the dynamics, especially above the main phase transition around 297 *K*. Interestingly, the Matryoshka model already proves to be sensitive enough to allow capturing differences between the gel and liquid phase in such lipid systems.

**Figure 5:**
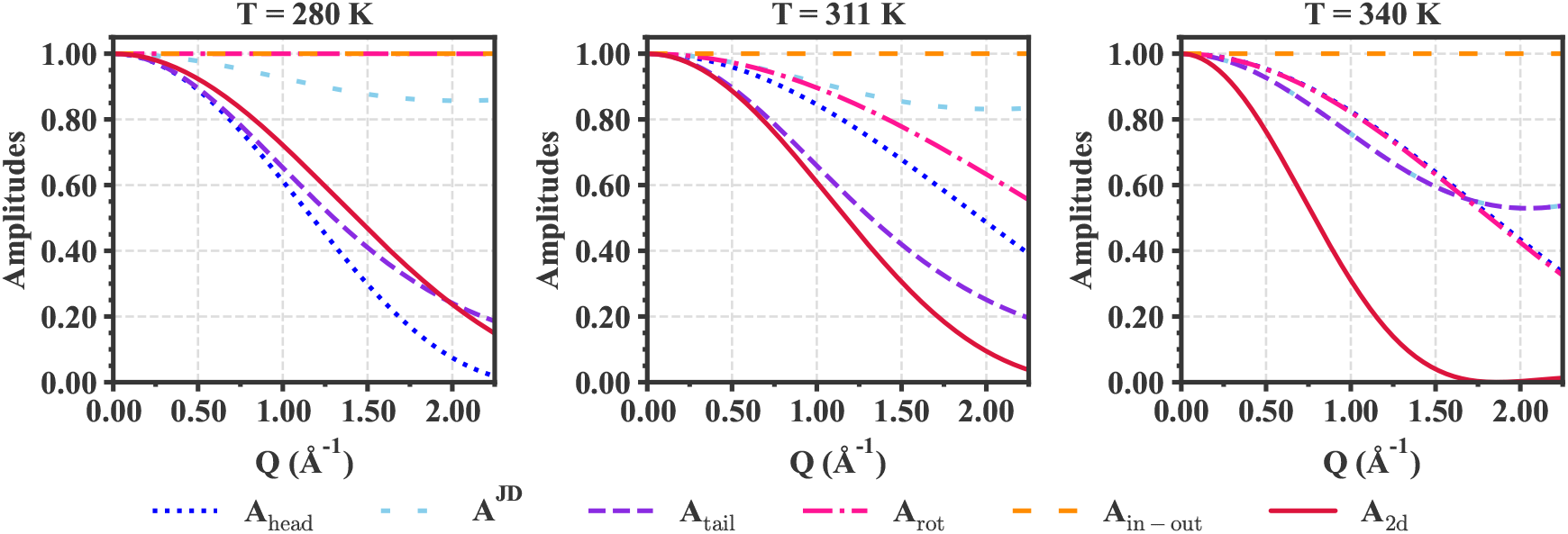
EISF or amplitudes of local motions (contributing to the 3-dynamical process) resolved in DMPC multilamellar bilayers at 135° (in-plane motions) as a function of *Q* for different temperatures. Lines correspond to expressions in Table 4 using parameters from Table 6. In terms of motion amplitudes, observable mouvements have amplitudes *A*_*i*_(*Q*) deviating from the horizontal line 1 as a function of *Q*. The deviation increases when *A*_*i*_(*Q*) decreases (i.e., the extent of motions increases) and vice versa, i.e., the area above the amplitude (*F*), 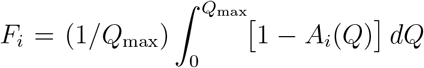, increases with the extension or amplitude of motions. More specifically,
  ⊳ *Fraction of dynamic H’s:* Values of *m* in Table 6 (given in %) are similar to that found in the literature [28], with *m* increasing with temperature.
  ⊳ *Individual motions:* The Debye-Waller factor is a gaussian function of *Q* (not shown in Fig. 5) with the mean-square displacements (very similar to that found in literature [28]) increasing with temperature (see Table 6). As a result, *F*_DW_(*Q*) increases with temperature.
  ⊳ *Internal motions*
    → *Backbone motions:* Backbone motions relate to movements of C-H groups described by the jump diffusion between two non-equivalent sites distant of *d*. As *d* is fixed (see Table 6), the change in the amplitude of backbone motions with temperature reflects the change in the probability *ϕ* of jump events that increases with temperature (see Table 6); note that *ϕ*(*T* = 340 *K*) ≈ 0.26 whereas *ϕ*_max_ = 0.5. Likewise, Fig. 5 shows that the amplitude of backbone motions increases as *F*_jump_(*Q*) increases with temperature.
    → *Tail motions:* Figure 5 shows that the diffusional motions of tails are clearly observables with almost same amplitudes for temperatures *T* = 280 *K* and *T* = 311 *K* as reflected by values of *R*_1_ in Table 6. And at *T* = 340 *K*, the *A*_*i*_(*Q*) of tail motions exhibit an increase and become similar to that of backbone motions. This behavior which does not go in the direction of the increase of parameters with temperature finds its origin in the imprecise value of *R*_1_ with large error bars (Table 6). It indicates possible variations of other parameters of the motion with temperature and / or relations between *R*_1_ and others parameters. Further analysis would be needed.
    → *Head motions:* Figure 5 shows that the amplitude of the head motions slightly increases with temperature at low temperature then decreases at higher temperature as reflected by changes in the head size *b*_H_ (see Table 6). Such a change of *b*_H_ at higher temperatures is interesting to note, especially since it could be indicative of an interference in the analysis between internal rotational motions of the heads around their axes and the molecule rotational movements of lipid molecules (see *Rotational motions* and Figure 5); both involving the H-atoms in the head group. More analysis would be needed. For example, analyzing amplitudes in ⊥ geometry, where **Q** is perpendicular to the membrane and only the movements of the heads are observable but not those of molecule rotation (see Table 4), could give informative indications on how to disentangle the motions.
  ⊳ *Molecule motions*
    → *Rotational motions:* It appears that the amplitude of the rotational motions of lipid molecules about lipid axis are weak (*A*_rot_(*Q*) ∼ 1) at low temperature and increases (*A*_rot_(*Q*) *<* 1) when the temperature increases thus resulting in an increase of the H-radius *R*_H_ of lipid molecules with temperature (see Table 6).
    → *In-out the plane motions:* The in-out of the plane motions of lipid molecules are not observable for the || geometry where the neutron scattering vector **Q** is parallel to the membrane (see Table 4). In this case, the amplitude remains the horizontal line, *F*_in*−*out_(*Q*) = 0, for all pairs of (*Q, T*) as shown in Fig. 5.
    → *In-plane 2d diffusion motions:* The in-plane diffusion of lipid molecules occurs within a cage of radius *R*_||_ formed by the neighboring lipid molecules. Figure 5 shows that the deviation *F*_2d_(*Q*) increases as the temperature gets higher, thus indicating that *R*_||_ increases with the temperature (see Table 6).

Finally, let us underline that some parameters of the model in the Table 6 are not comparable with structure parameters which could have been obtained from diffraction experiments, for example. Indeed, unlike the structural parameters, the model parameters relate to the local movements of the H atoms around their equilibrium positions. By definition, these dynamical parameters are different and will generally be smaller than their structural counterparts. For example, the values of the H-radius, *b*_H_, of the head group are slightly smaller than the experimental data (∼ 2Å) of the head size [26, 29]. Likewise, the values of the smallest radius *R*_1_ for the tail motions are smaller than the experimental values (∼ 2.2Å) of chain spacings at the start of the tails [26, 29]. As for the H-radius, *R*_H_, it can be compared to the span of the lipid molecule given by *D*_H1_ sin(*α*), where *D*_H1_ (similar but different from *d*_H_ in Fig. 2) is the size of the head group and *α* the tilt angle of the head. We find that *R*_H_ is smaller than the lipid span for *α* given in Table 6 and *D*_H1_ ∼ 5Å obtained from diffraction [26, 29].

## 4. Concluding summary

Our main motivation in developing this work has been to construct a framework for studying and describing local dynamics of lipid molecules in membrane layers. The main results of this work can be summarized as follows:

- We have developed the dynamical Matryoshka model which describes the local dynamics of lipid molecules in a membrane layer as a nested hierarchical convolution of three motional processes (see, Table 1):
  i. individual motions described by vibrational movements of H-atoms;
  ii. internal motions including the motions of the lipid backbone described by a jump dynamics between two non-equivalent sites, the head motions described by a rotational diffusion about the head axis plus a flip-flop or jump dynamics of the head tilted angle about the lipid axis, and the motions of the tail described by two-dimensional diffusions of H-atoms within circles of increasing radii along the tails;
  iii. molecule motions of the lipid molecule as a whole including the rotational diffusion about the lipid axis, in-out of the plane motion and the in-plane local diffusion of the lipid molecule within a cage.
  The model includes seven motions in total (see, Table 2) with the analytical expressions of all associated ISFs provided in Table A.7.
- For the purpose of analyzing QENS experimental data, we have derived an analytical expression for the aggregated ISF of the Matryoshka model which involves an elastic term plus three inelastic terms of well-separated time scales. In doing so, we obtain the relationships between the amplitudes and rates of the aggregated ISF and those of the local movements of the initial model (see, Table 3).
- As a check of the model, we have shown that the theoretical aggregated ISF fit very well the QENS experimental data on a DMPC sample and allow extracting the dynamical parameters of the model.

Although we have shown that the development outlined above already describes very well QENS experiments, it can be complemented with analyzes including both the amplitudes, *A*_*i*_(*Q*), and rates, Γ_*i*_(*Q*), for various samples of lipid systems in different geometries. Such work is carried out in [21, 22]. The Matryoshka model provides a framework within which several improvements can be carried out if necessary. For example, for local motions, we have described the motions of the backbone by a jump dynamics between two sites. It is quite possible to replace it with other movements (e.g., jump dynamics between three sites) that the associated EISF (which contributes in the amplitudes of intermediate and fast motions) does not cancel at large *Q*, as experimental evidence indicates that the movements of the backbone should be spatially discrete. The Matryoshka model outlined above can be further extended to include in the same setting long range jumps for in-plane lipid diffusion and collective undulation modes of lipids. In terms of the time scales discussed here, these motions belong to the class of slow or very slow motions (Γ ∼ 1 *µ*eV). Such a work is in progress. Finally, to be complete in the analysis of QENS data, it might turn out necessary to develop models predicting how the parameters of local motions would change with temperature, for example.

## Acknowledgements

The authors thank the Institut Laue-Langevin for the allocation of the beam time to perform the experiments. We also thank Francesca Natali for her help on the experimental aspects of QENS. AC is supported by the JP Aguilar scholarship from Fondation CFM for her PhD thesis.

## Data Availability

The datasets generated and analyzed during the current study are available from the corresponding author on reasonable request.

## Author Contributions

Dominique J. Bicout: Conceptualization, Methodology, Writing - Original Draft, Review & Editing, Supervision. Aline Cisse: Software, Validation, Formal Analysis, Writing - Review & Editing. Tatsuhito Matsuo: Software, Validation, Formal Analysis, Writing - Review & Editing. Judith Peters: Investigation, Writing - Review & Editing, Supervision, Funding acquisition.

## Competing Interests

The authors declare that they have no financial or non-financial competing interests.

## Appendix A. Derivations of expressions of ISF

The key quantity in neutron scattering is the incoherent structure function (ISF), defined as,

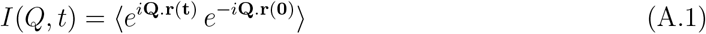

where **Q** is the neutron scattering wave vector, **r**(**t**) and **r**(**0**) the hydrogen atom positions at *t* = 0 and time *t*, respectively, and the sign ⟨…⟩ represents an ensemble average over all positions **r**(**t**) and **r**(**0**) in the potential of mean force, *V* (**r**). The mathematical expressions of all the ISFs considered in this work are gathered in Table A.7. For the ISFs whose expressions had already been derived elsewhere, we have indicated the associated references, and for the others we derive the expressions in this Appendix. This will be the case for head rotational movements with flipflop and the molecular in-out-of the plane diffusion of the lipid molecule in a harmonic potential.

**Table A.7:**
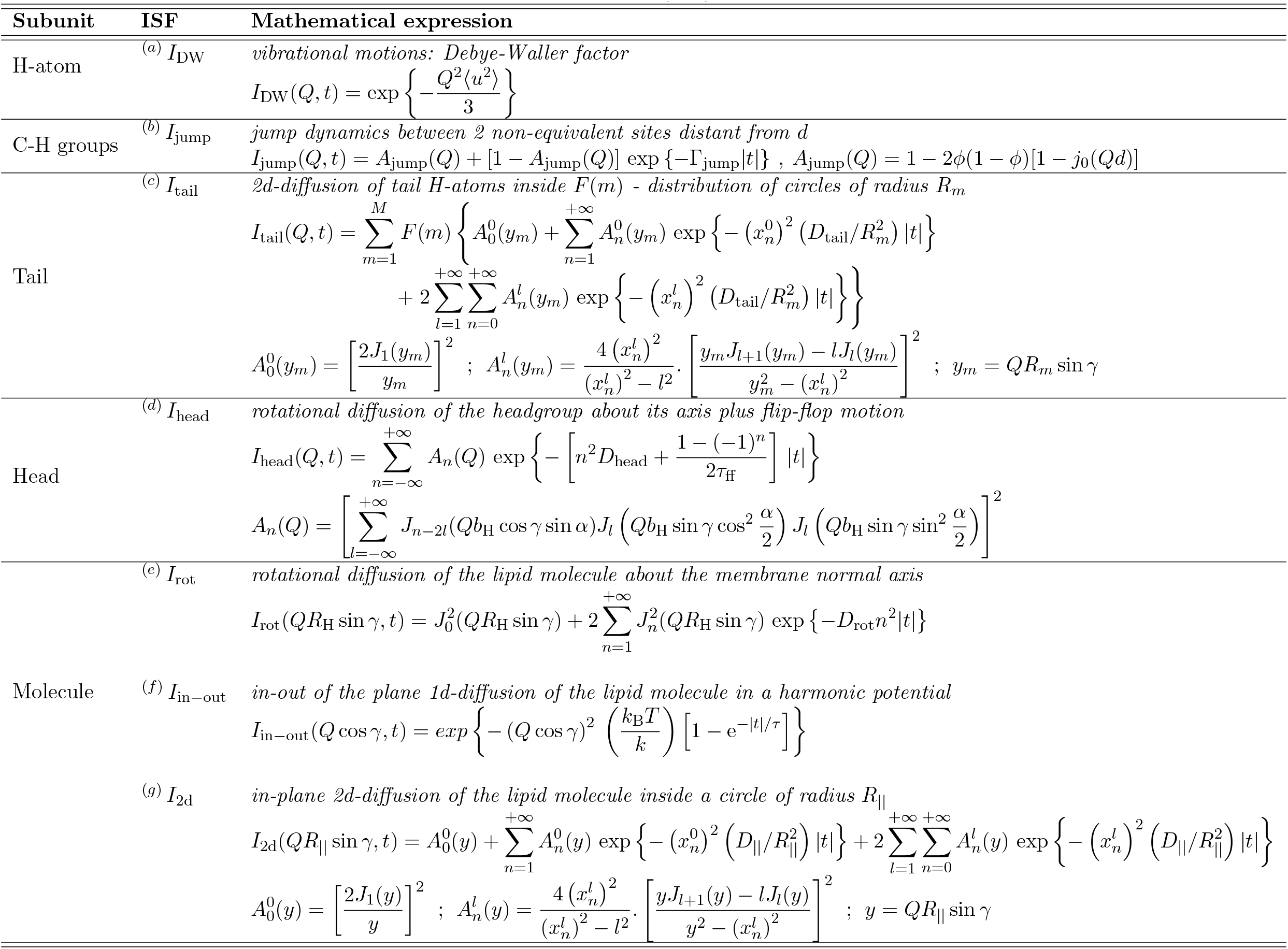
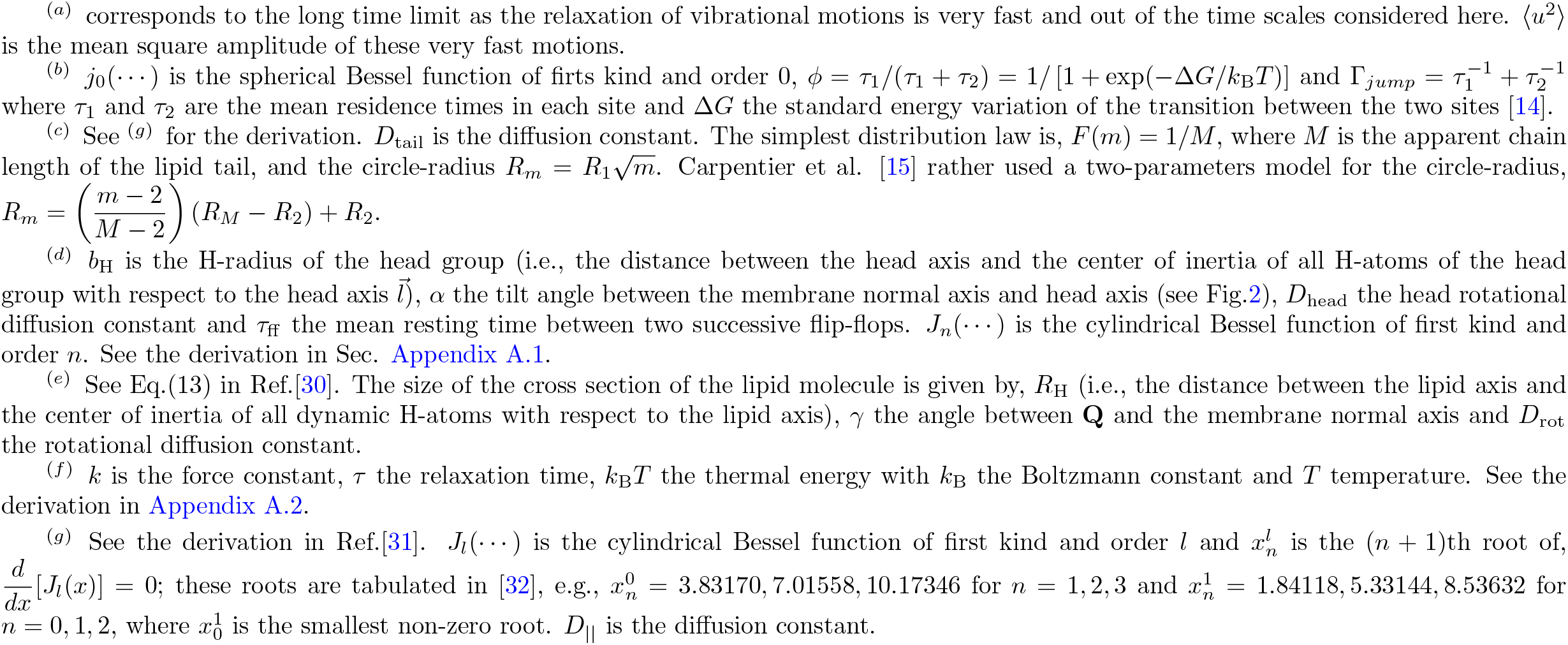
Incoherent Structure Functions (ISF) of dynamical processes.

### Appendix A.1. Rotational diffusions and flip-flop of headgroup

We will consider the head group as an ellipsoid of circular cross-section of diameter 2*b*_H_ and directing vector 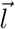 (see Fig.2), where *b*_H_ is the distance between the head axis and the center of inertia of all H-atoms of the head group with respect to the head axis 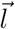 and 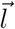 is given in the orthonormal frame 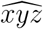 of basis 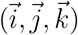 by,

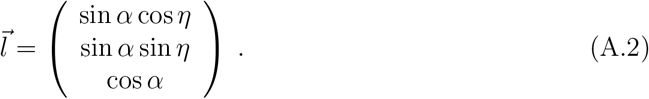

The angles are defined in Fig.A.6.

**Figure A.6:**
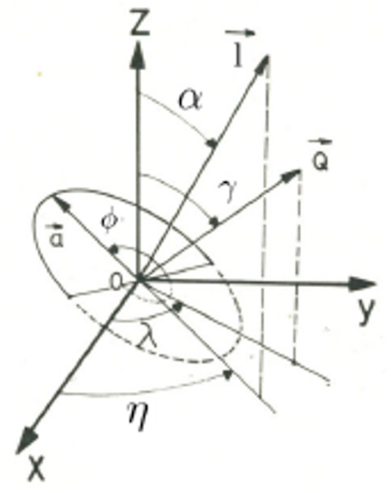
Definition of angles for the rotations about 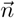 (parallell to *z*) and 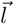 directions and the flip-flop motion of 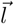 about 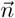. Adaptation from [30].

That thus makes it possible to define a local orthonormal coordinate system 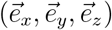 associated with the head group as follows,

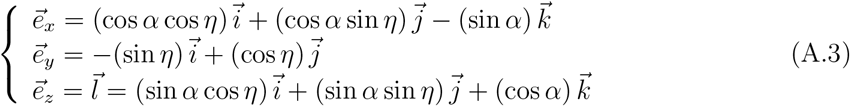

from which the matrix of passage from 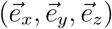 to 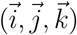 can be determined as,

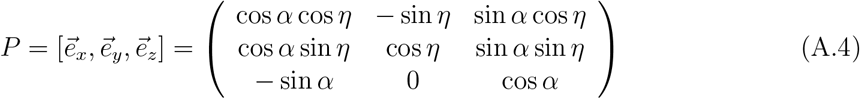

Note that *P* = *R*_*z*_(*η*)*R*_*y*_(*α*) results from the operation of two rotations where *R*_*y*_(*α*) describes the rotation of an angle *α* about *y*−axis and *R*_*z*_(*η*) the rotation of an angle *η* about *z*−axis. The head group position vector 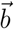 in the local head group frame 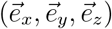 is given by,

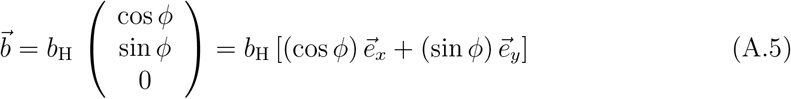

where *ϕ* is the angle between 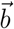 and 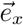. Thus, the head group position vector **r** in the frame 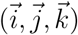 is obtained using the passage matrix as,

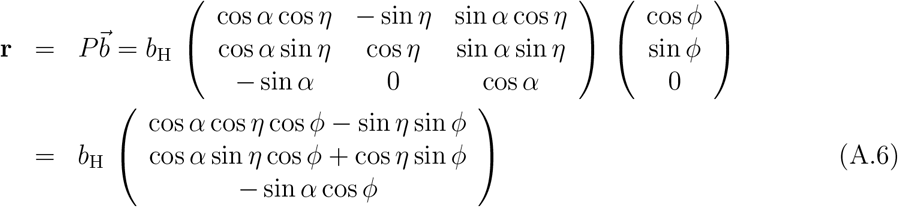

Note that, 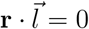 as expected since 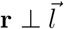.

Let the *z*−axis be the normal to the membrane, the scattering wave vector **Q** is given by,

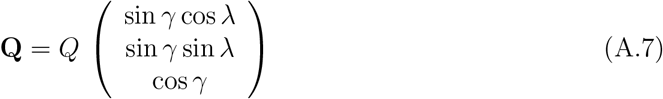

where *γ* is the angle between 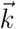 and **Q** and *λ* the angle between the projection of **Q** onto the *x* − *y* plan and 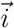. Now, the scalar product entering in the calculation of the ISF writes as,

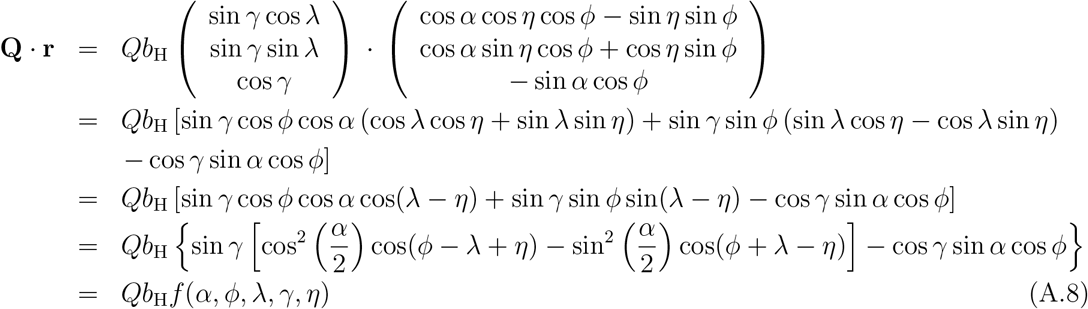

The head group performs two independent motions:

- Uniaxial rotational diffusion of angle *ϕ* about the 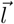 –axis and diffusion constant *D*_head_ described by the Green’s function:

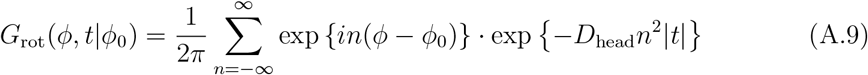

with the equilibrium distribution, 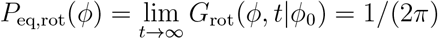,
- Flip-flop motion of the 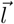−axis between angles *θ* = −*α* and *θ* = *α* and residence time *τ*_ff_ described by the jump diffusion with the Green’s function:

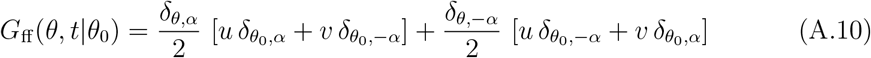

where,

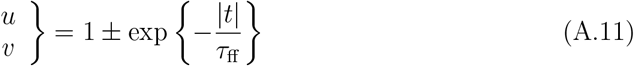

with the equilibrium distribution, 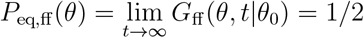.

For experimental configurations where the direction of the scattering vector **Q** with respect to the membrane normal is set constant (i.e., the angle *γ* is set constant for all *λ*), the ISF is calculated as,

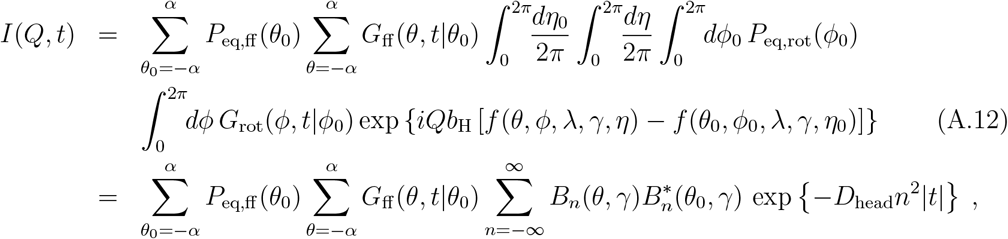

where,

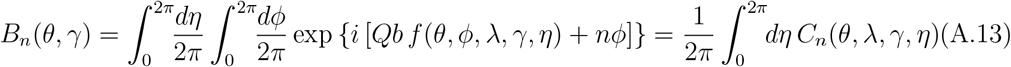

Using G_ff_(*θ, t*|*θ*_0_) in Eq.(A.10) back into *I*(*Q, t*), we have:

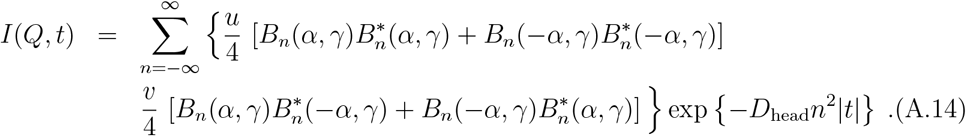

We have,

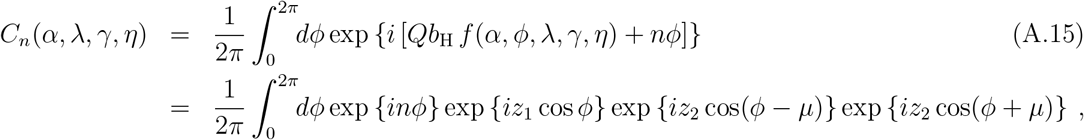

where,

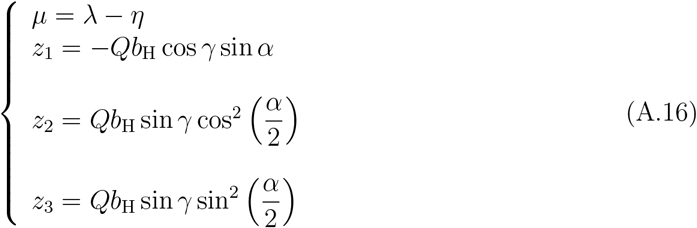

Now, using the following relations,

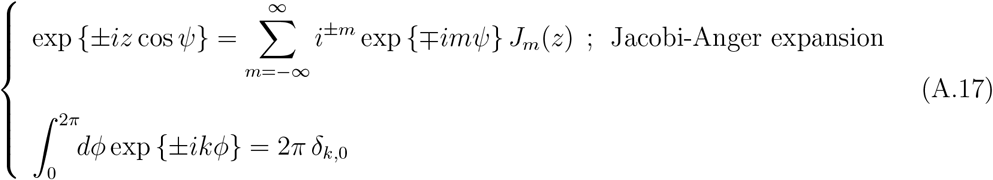

back into Eq;(A.15), we obtain:

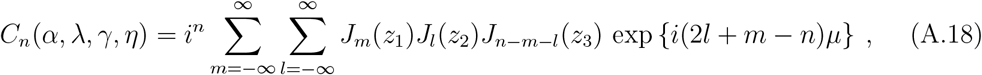

where *J*_*m*_(···) is the Bessel function of first kind and order *m*. It follows that,

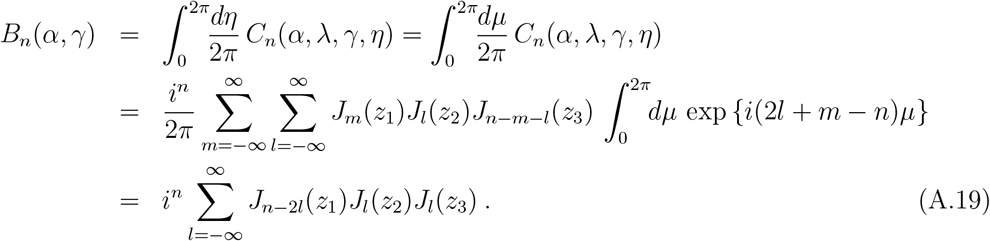

Using the relation for the Bessel functions, *J*_*n*_(− *x*) = (− 1)^*n*^*J*_*n*_(*x*), we obtain the relation: *B*_*n*_(*α, γ*) = (−1)^*n*^*B*_*n*_(−*α, γ*). Now, using this relation relation in Eq.(A.14), we finally obtain,

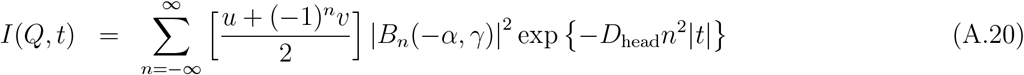

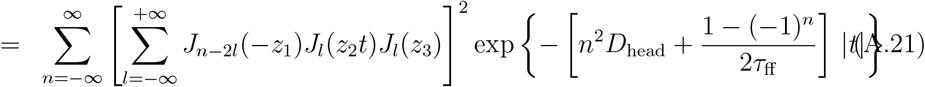

### Appendix A.2. 1d Diffusion in a Harmonic Potential

Let the *z*–axis coincides with the normal to the membrane, we consider that each lipid molecule, of coordinate *z*, is undergoing as a whole a 1d diffusion parallel to *z*–axis in a harmonic potential of mean force, *V* (*z*) = *kz*^2^*/*2, of force constant *k* and relaxation time *τ*. The Green’s function, *G*(*z, t*|*z*_0_), describing such motions for the lipid molecule is given by,

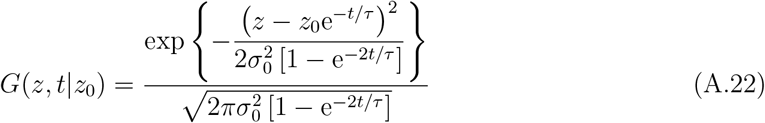

with the equilibrium distribution of lipid molecule positions given by,

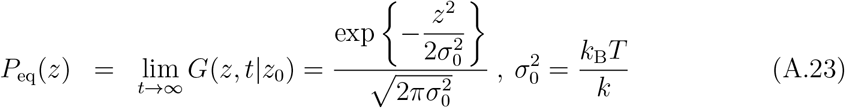

Denoting by *γ* the angle between the *z*–axis and the scattering **Q**, the ISF can be written as,

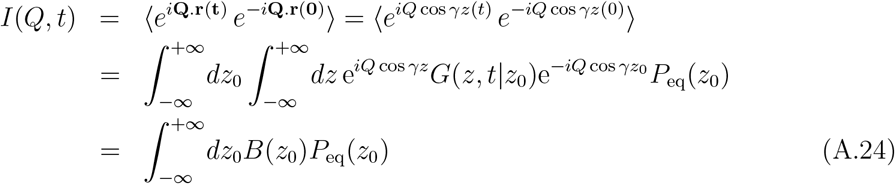

where,

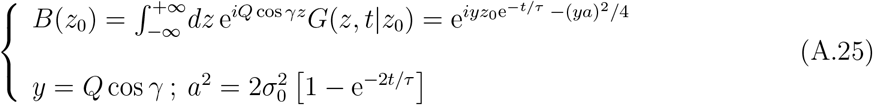

Then,

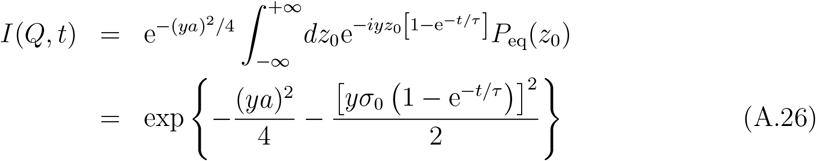

Finally, we obtain:

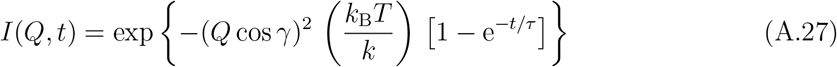

## Appendix B. Derivation of aggregated expressions of *B*’s amplitudes and Γ’s rates

Starting from a time dependent ISF, the aim of the aggregation is in general to derive an approximate expression of ISF as an expansion of only few relaxation functions, *E*_*k*_(*Q, t*) such that, *E*_*k*_(*Q, t* = 0) = 1 and *E*_*k*_(*Q, t* → ∞) = 0. For our purpose, we deal with single exponential relaxation function, *E*_*k*_(*Q, t*) = exp {−Γ_*k*_*t*}, where Γ_*k*_(*Q*) is the relaxation rate. The ISF to derive an aggregation is given in the right-hand-side of Eq.(10) as,

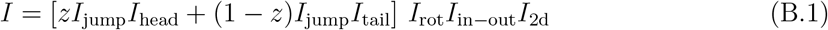

where *I*_*α*_(*Q, t*) are the ISF’s of local motions described in the Matryoshka model and given as [16],

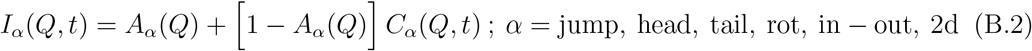

where *A*_*α*_(*Q*) is the amplitude and *C*_*α*_(*Q, t*) the relaxation function such that, *C*_*α*_(*Q, t* = 0) = 1 and *C*_*α*_(*Q, t* → ∞) = 0. As, in general, *C*_*α*_(*Q, t*) is a multi-exponential function of time with decay rates functions of *Q*, the first step toward the aggregation is to derive a single exponential approximation of *C*_*α*_(*Q, t*) as [16],

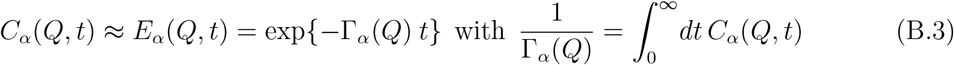

where Γ_*α*_(*Q*) is the *Q*–dependent relaxation rate. Next, the goal is to derive the amplitudes and relaxation rates such that Eq.(B.1) can be rewritten as,

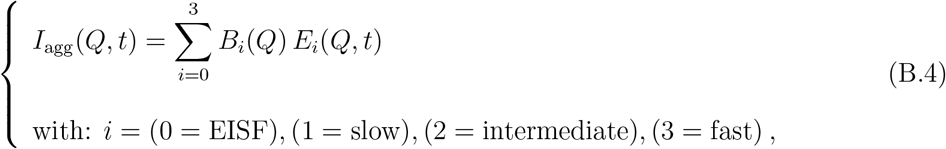

where *B*_*i*_(*Q*) is the amplitude and *E*_*i*_(*Q, t*) the relaxation function with relaxation rates such that, Γ_0_ = 0 *<* Γ_slow_ *<* Γ_intermediate_ *<* Γ_fast_. To proceed, expanding Eq.(B.1) yields,

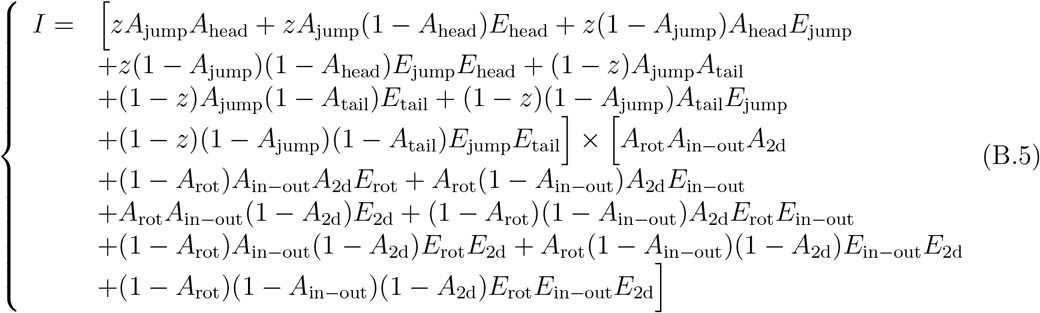

Next, we use the following rules,

- Hierarchy of relaxations: the hierarchy of relaxation time scales of motional processes in the Matryoshka model indicates that: Γ_jump_ ∼ Γ_tail_ *>* Γ_head_ ∼ Γ_rot_ *>* Γ_in*−*out_ ∼ Γ_2d_, implying that, *E*_jump_ ∼ *E*_tail_ *< E*_head_ ∼ *E*_rot_ *< E*_in*−*out_ ∼ *E*_2d_ for all *t*;
- Operation rules: for timescales Γ_*i*_*t* ∼ 1, we have:
  ⊳ Summation: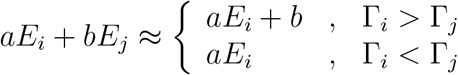
  ⊳ Product: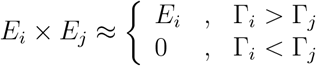

to rearrange Eq.(B.5) into 4 terms and derive the aggregated approximation in Eq.(B.4) where,

- **EISF:** collects all time independent terms in Eq.(B.5). Taking the *t* → ∞ limit (i.e., setting all *E*_*α*_(*Q, t* → ∞) = 0) in Eq.(B.5) gives,

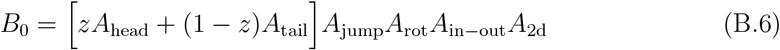
- **Slow motions:** collect all terms involving *E*_in*−*out_ and *E*_2d_ in Eq.(B.5).

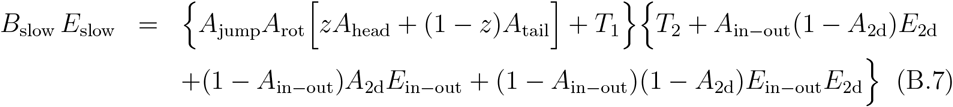

in which *T*_1_ groups terms involving *E*_jump_, *E*_tail_ and *E*_head_ and *T*_2_ terms involving the products of *E*_rot_ with *E*_in*−*out_ and *E*_2d_ (i.e., terms ≈ *E*_rot_). For timescales Γ_2d_*t* ∼ Γ_in*−*out_*t* ∼ 1, both *T*_1_ and *T*_2_ vanish as they relax faster to zero and, therefore, will be omitted. Now, taking the *t* → 0 limit in Eq.(B.7) (without *T*_1_ and *T*_2_) gives,

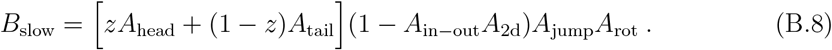 And, taking the *t* → 0 limit in the time derivative of Eq.(B.7) gives,

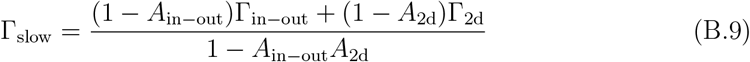
- **Intermediate motions:** collect all terms involving *E*_head_ and *E*_rot_ in Eq.(B.5). For timescales Γ_head_*t* ∼ Γ_rot_*t* ∼ 1, we use the rules above, *E*_in*−*out_ ∼ *E*_2d_ ≈ 1 and *E*_jump_ ∼ *E*_tail_ ≈ 0 to first reduces Eq.(B.5) to,

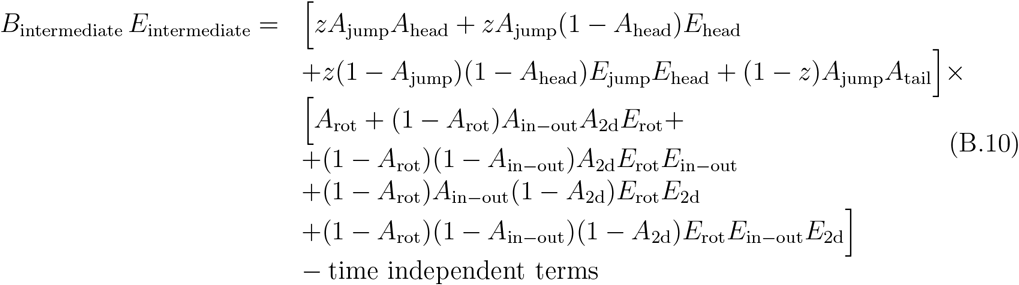

and next, *E*_rot_*E*_in*−*out_ ≈ *E*_rot_, *E*_rot_*E*_2d_ ≈ *E*_rot_, *E*_rot_*E*_in*−*out_*E*_2d_ ≈ *E*_rot_ and *E*_jump_*E*_head_ ≈ 0 to,

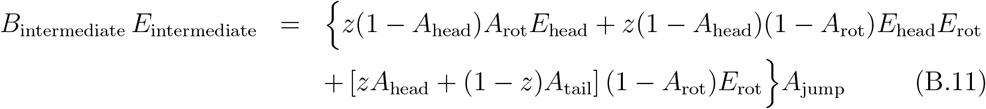 Taking the *t* → 0 limit in Eq.(B.11) gives,

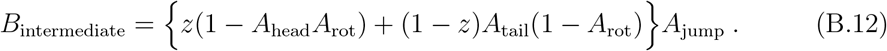

And, taking the *t* → 0 limit in the time derivative of Eq.(B.11) gives,

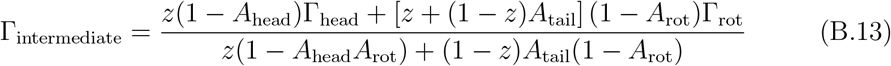
- **Fast motions:** collect all terms involving *E*_jump_ and *E*_tail_ in Eq.(B.5). For timescales Γ_jump_*t* ∼ Γ_tail_*t* ∼ 1, we use the rules above, *E*_in*−*out_ ∼ *E*_2d_ ∼ *E*_rot_ ∼ *E*_head_ ≈ 1 to reduces Eq.(B.5) to,

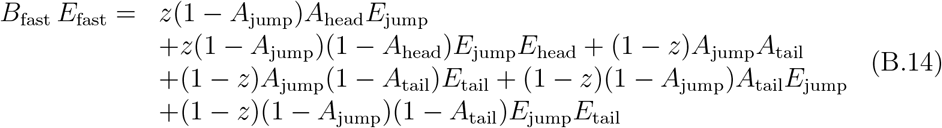

or,

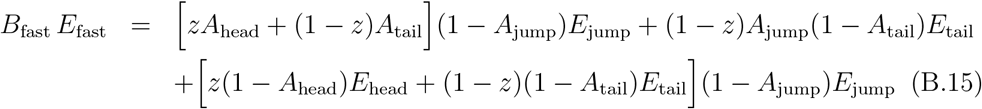

Taking the *t* → 0 limit in Eq.(B.15) gives,

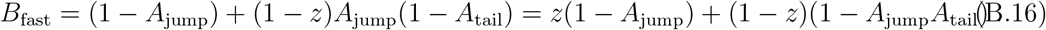

And, taking the *t* → 0 limit in the time derivative of Eq.(B.15) gives,

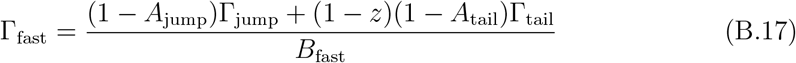

## Appendix C. Relation between experimental *A*’s and theoretical *B*’s amplitudes

Given that the theoretical aggregated ISF in Eq.(B.4) writes as follows,

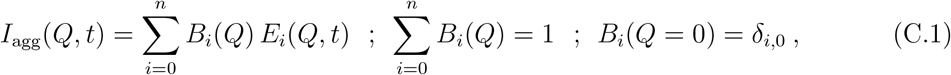

we assume that the experimental ISF can also be written in the same way as,

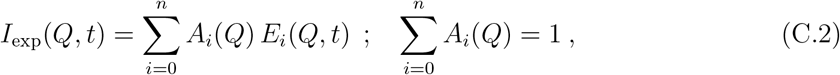

with the same *E*_*i*_(*Q, t*) but different amplitudes *A*_*i*_(*Q*) and such that, 0 *< A*_*i*_(*Q* = 0) *<* 1, ∀*i*. The correspondence between *I*_agg_(*Q, t*) and *I*_exp_(*Q, t*) can be written as,

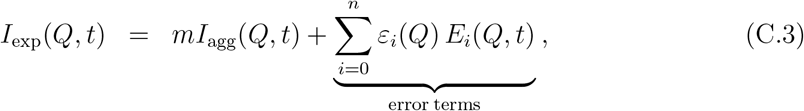

where *m* is the fraction of observable mobile H-atoms and *ε*_*i*_(*Q*)’s the errors accounting for the fraction of immobile H-atoms and other experimental errors like multiple scatterings, etc. Collecting in Eq.(C.3) terms under the same relaxation function *E*_*i*_(*Q, t*), we obtain the general relationship between *B*(*Q*)’s and *A*(*Q*)’s as,

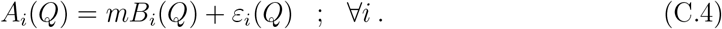

From the point of view of the analysis of experimental data, the expression in Eq.(C.4) involves *n* + 2 unknowns to determine: *m* and *n* + 1 functions *ε*_*i*_(*Q*). Therefore, in the absence of any information we use the closure relation (obtained by construction) satisfied by the unknowns and assume that the errors of the quasi-elastic terms (*i >* 0) are all identical, i.e.,

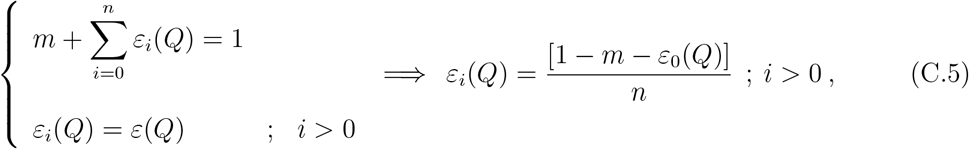

thus reducing the number of unknowns from *n* + 2 to 2. Using this back in Eq.(C.4), we obtain,

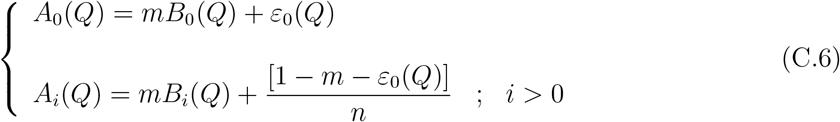

where the remaining unknowns are *m* and *ε*_0_(*Q*).

## Notes

### Competing Interest Statement

The authors have declared no competing interest.

